# Hepatitis B virus proteome analysis identifies apolipoprotein C1 facilitating particle production and virus entry

**DOI:** 10.64898/2026.04.03.716119

**Authors:** Shangqing Yang, Firat Nebioglu, Minh Tu Pham, Yen-Chia Lin, Andreas Pichlmair, Shirin Nkongolo, Pietro Scaturro, Stephan Urban, Stefan Seitz, Ralf Bartenschlager

**Author notes:** Contributed equally to this work (Shangqing Yang, Firat Nebioglu). **Correspondence:** Prof. Dr. Dr. h.c. Ralf Bartenschlager, Department of Infectious Diseases, Molecular Virology, Center for Integrative Infectious Disease Research (CIID), University of Heidelberg, Im Neuenheimer Feld 344, 1st. Floor, 69120, Heidelberg, Germany.

## Abstract

**Background & Aims:** Antiviral therapies targeting hepatitis B virus (HBV) suppress viral replication, but rarely achieve functional cure. Understanding HBV-host cell interaction is crucial for developing novel therapeutic approaches. Here, we report host cell proteins associated with HBV virions and filamentous subviral particles (fSVPs) and characterize one of them, apolipoprotein C1 (ApoC1), mechanistically.

**Methods:** Highly purified HBV virions and fSVPs were obtained by sequential use of several biophysical methods. Particles were analyzed by mass spectrometry and associated proteins were evaluated phenotypically using an HBV infection model. The top hit, ApoC1 was characterized in detail.

**Results:** Associated with virions and fSVPs, we identified in addition to known chaperones such as HSP90AB1 and HSC70, several apolipoprotein-related factors. RNAi-based phenotypic validation identified strongest effects for ApoC1, likely due to two complementary effects. First, ApoC1 depletion reduced intracellular cholesterol level impairing HBV infection and SVP production, which was compensated by exogenous cholesterol substitution. Second, ApoC1 that is mainly enriched in high-density lipoprotein (HDL), associates with HBV virions and fSVPs and increases HBV infectivity. The same was found for hepatitis D virus (HDV), a satellite virus utilizing HBV envelopes. Supplementation of exogenous HDL enhanced infection most likely via scavenger receptor class B type 1 (SR-B1), the natural HDL receptor. Consistently, inhibition of SR-B1 suppressed HBV and HDV infection.

**Conclusions:** We established a method for obtaining highly purified HBV virions and fSVPs and identified the HDL component ApoC1 to associate with both particle types. ApoC1 promotes HBV and HDV infection most likely via SR-B1 facilitating viral entry.

## Introduction

Chronic hepatitis B virus (HBV) infection is a major public health burden affecting an estimated 254 million people worldwide^1^. They have a high risk to develop severe liver diseases, including liver cirrhosis and hepatocellular carcinoma (HCC), accounting for a total of 1.1 million deaths per year^1^. Antiviral therapies, such as the use of nucleot(s)ide analogues, are rarely curative and require lifelong treatment. These drugs suppress viral replication but do not eliminate the covalently closed circular (ccc) DNA, the viral persistence reservoir residing in the nucleus of infected cells^2, 3^. In addition, excessive amounts of non-infectious subviral particles (SVPs) are released from infected hepatocytes, causing dysfunction of the immune response targeting the major viral antigen, the hepatitis B surface antigen (HBsAg). Therefore, to achieve curative therapy for chronic hepatitis B, novel approaches are urgently needed.

HBV particles possess a lipid envelope containing 3 viral glycoproteins called S, M and L (small, medium and large, respectively). They share the same C-terminal S-region but differ by having additional N-terminal extensions: the preS2 region of M and the preS1 region of L, the latter binding to the viral entry receptor NTCP (sodium taurocholate co-transporting polypeptide)^4^. Initial attachment of HBV to hepatocytes is facilitated by heparan sulfate proteoglycans (HSPGs) interacting with the viral surface proteins^5^. Subsequently, the preS1 region specifically binds to NTCP, followed by virion entry and nucleocapsid release into the cytoplasm. Upon transport into the nucleus, the relaxed circular DNA (rcDNA) is converted into the cccDNA by the host DNA repair system^6, 7^. cccDNA associated with cellular histones forms a stable episome from which all HBV RNAs are transcribed^8^. Transcription requires the action of the HBx protein preventing the epigenetic silencing of the cccDNA template^9, 10^. Infected cells release an array of HBV-derived particles, including infectious virions (Dane particles), non-enveloped nucleocapsids with and without viral RNA and excessive quantities of non-infectious SVPs, the latter contributing to selective immune tolerance^11, 12^. Depending on the L-M-S ratio, SVPs can have two morphologies: spherical SVPs (sSVPs), 20-25 nm in diameter, predominantly composed of S and released via the constitutive secretory pathway; filamentous SVPs (fSVPs) containing S and also L and released from cells via the ESCRT/MVB pathway^13^. The latter is also used for HBV virion release.

Several studies reported the incorporation of host cell proteins into HBV nucleocapsids or virus particles. Early work demonstrated that the interaction between HBV polymerase and pregenomic (pg) RNA requires the heat shock protein HSP90, detected in purified nucleocapsids together with the chaperone HSP70^14^. A more recent study reported that DNase1 is upregulated and encapsidated under hypoxic conditions, resulting in degradation of HBV DNA within secreted virions^15^. In addition, cyclophilin A (PPIA) and apolipoprotein E (ApoE) have been reported to be associated with secreted particles^16, 17^. Consistent with the latter, the reverse cholesterol transport pathway in hepatic macrophages was found to be hijacked by HBV to facilitate virion uptake by hepatocytes^18^.

Despite increased understanding of the HBV replication cycle, important gaps exist. One is the lack of knowledge of host cell proteins carried along within HBV virions and delivered into the infected cell. Here, we established a particle purification pipeline enabling the proteomic analysis of HBV virions and fSVPs with high specificity and sensitivity. We identified apolipoprotein C1 (ApoC1), a component of high-density lipoprotein (HDL), as a critical host cell dependency factor and reveal its contribution to HBV infection and particle production via a dual-mode of action.

## Materials and methods

### HBV particles production and purification

HepAD38 cells containing an overlength HBV genome under the control of a tetracycline-inducible promoter were cultured as previously described^19^. HBV production was induced by omitting doxycycline from the culture medium. Culture supernatants were collected every four days, sterile filtered and stored at 4℃ until further processing. HBV particles were purified as described in the Results section and Supplementary Materials and Methods.

### HBV and HDV infection

One day after seeding, HepG2-NTCP A3 cells were infected with HBV as described in the results section. For HDV infection, Huh7-NTCP cells were inoculated with a multiplicity of infection (MOI) of 5 HDV GE per cell in the presence of 4% PEG 8000 and 1.5% DMSO for 24 h. Viral replication was determined as reported in supplementary materials and methods.

### Antibody neutralization assay

Purified HBV stocks were incubated with various concentrations of ApoC1-specific antibody for 1 h at room temperature, then added to HepG2-NTCP A3 cells in the absence of PEG 8000. After 24 h, cells were washed with PBS and cultured for additional seven days. Secreted HBsAg, HBeAg and HBV DNA levels were determined by ELISA and qPCR, respectively, while cell lysates were used to measure HBcAg abundance by western blot.

### Ethics statement

Use of human clinical samples was approved by the Ethics Committee of the Medical Faculty of Heidelberg University (approval no. S-522/2024) and conducted in accordance with the Declaration of Helsinki. Written informed consent was obtained from all patients whose samples were collected prospectively. For retrospectively collected samples, the requirement for informed consent was waived by the ethics committee.

For all further details, see supplementary materials and methods.

## Results

### Purification of HBV virions and filamentous SVPs

To obtain HBV virions and fSVPs, we used two HBV-containing HepG2-derived cell lines (Fig. 1A). First, HepG2-HB2.7 cells containing only a partial 2.7 kb HBV genome encoding the envelope proteins and the HBx protein and supporting only SVP production. Second, HepAD38 cells harboring a 1.1-overlength HBV genome under a tet-off inducible promoter and releasing both HBV virions and SVPs upon tetracycline removal from the culture medium^19^. The parental HepG2 cell line was used as control to determine contaminating proteins in the proteomics analysis (Fig. 1A). By continuously measuring HBsAg and HBeAg levels in the supernatant of long-term cell cultures (2-3 months), we monitored HBV particle production. Since we anticipated contamination of the preparations by components present in fetal bovine serum (FBS) used for cell cultivation, we generated 5 independent biological replicates, three from cells cultured in the presence of 10% FBS (replicates 1-3) and 2 preparations with cells cultured in medium containing 5% FBS (replicates 4-5).

**Fig. 1.**
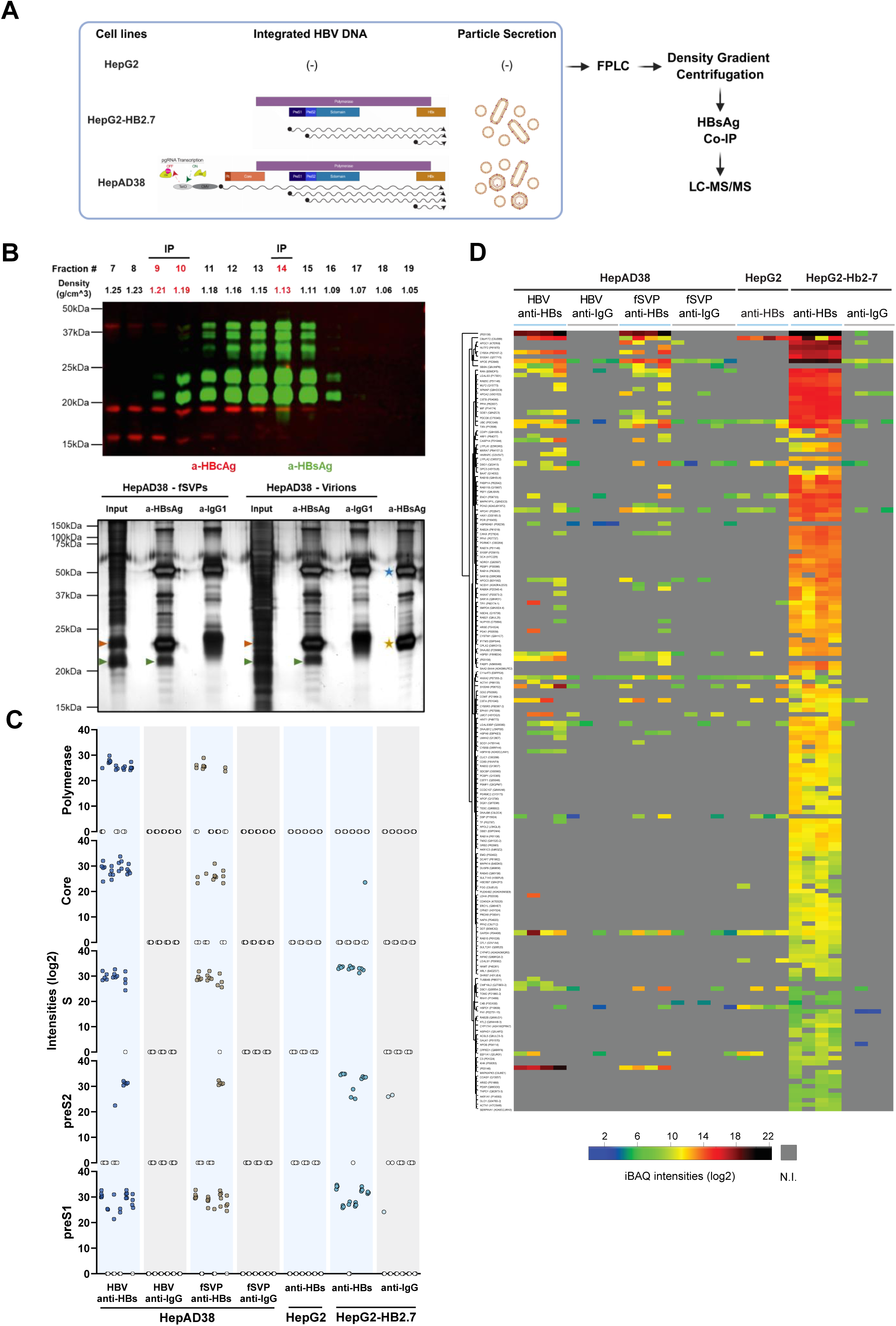
Purification of HBV virions and filamentous SVPs. (A) Purification pipeline of HBV virions and fSVPs from HepG2-HB2.7 and HepAD38 culture supernatants. (B) Upper panel: Sucrose fractions of one representative equilibrium density gradient were analyzed by dual-fluorescence western blot detecting HBV envelopes (green) and core proteins (red), with densities specified above each lane. Fractions enriched for virions (9-10) or fSVPs (14) were used for HBsAg-specific pull-down. Lower panel: Pull-down samples and corresponding inputs were analyzed by SDS-PAGE and silver staining. Arrowheads indicate glycosylated (orange) and non-glycosylated (green) small HBs proteins; stars indicate IgG heavy (blue) and light (gold) chains. (C) Viral peptides identified by AP-LC-MS/MS across all 5 biological replicates are categorized according to viral proteins. Individual values represent spectral intensities (log2) of unique peptides from pull-down samples across independent biological replicates. (White circles = not identified). (D) Proteomic analysis of affinity-purified particles by AP-LC-MS/MS (n≥4). The heatmap represents intensity-based quantification (iBAQ) values of significantly enriched proteins in virion or fSVPs pull-downs (log2). Each column corresponds to one pull-down sample from one independent biological replicate. N.I.= not identified.

To obtain mass spectrometry-grade HBV particles, we employed our previously reported purification protocol comprising FPLC with heparin column and size exclusion chromatography followed by density gradient centrifugation (Fig. 1A)^20^. To remove the majority of spherical SVPs we intentionally selected only the early fractions of the size-exclusion chromatography, which contain larger particles such as Dane particles and filamentous SVPs. Dual fluorescence western blot analysis of the fractions collected after ultra-centrifugation demonstrated distinct distribution of particles: naked capsids (density ∼1.23 g/cm^3^; indicated also by HBeAg that cross-reacts with core protein); virions (∼1.20 g/cm^3^; indicated by HBsAg and HBV DNA) and fSVPs (density of ∼1.13 g/cm^3^; indicated by high amounts of HBV envelope proteins and low HBcAg and HBV DNA) (Fig. 1B, upper panel and Fig. S1). Upon pooling of the virion fractions, they were analyzed for total protein, along with the selected fSVP fraction, by using silver staining. We observed high amounts of protein in both samples, dampening detection sensitivity of the HBV proteome (Fig. 1B, lower panel, input lanes; for gradient fractions see Fig. S2A). The same was true for fSVPs purified from culture supernatant of HepG2-HB2.7 cells (for gradient fractions see Fig. S2B and S3A, input lane). To reduce impurities, we performed HBs-specific immunoprecipitation (IP); IP with an IgG isotype was used as specificity control. Protein amounts and complexities of the samples were greatly reduced after IP for both the virion and fSVP fractions derived from HepAD38 cells and for fSVPs derived from HepG2-HB2.7 cells (Fig. 1B, lower panel and Fig. S3A, respectively). After IP, HBc protein remained detectable in the virion fractions indicating that integrity of the particles was retained (Fig. S3B, left panel). As expected, re-probing the blots with HBs-specific antibodies revealed HBV envelope proteins of different isoforms in the virion fractions (Fig. S3B, right panel).

### Proteomics analysis of purified HBV virions and filamentous SVPs

Proteins contained in IP-purified virion and fSVP samples were identified by liquid chromatography-tandem mass spectrometry (LC-MS/MS). For each experimental condition, four or five biologically independent replicates of affinity-purified complexes obtained from IgG or HBs-specific immunoprecipitations were collected and analyzed by quantitative mass-spectrometry. Although abundance of each unique viral peptide varied between the different replicates, HBV envelope proteins, core- and polymerase (P)-protein were consistently detected in all virion fractions with the best peptide coverage in replicate #5 (Fig. 1C and Table S1). Notably, we detected 5 different peptides derived from the HBV P-protein that is present in only one copy per particle^21^. In addition, we detected core- and envelope-specific peptides, whereas HBx- and HBe-specific peptides were not identified. In contrast, only some P- and much lower amounts of core-specific peptides were detected in the fSVP fraction from HepAD38 cells, demonstrating a strong depletion of virions from the fSVP fraction. In the case of fSVP fractions derived from HepG2-HB2.7 cell supernatants, much stronger signals for HBV envelope proteins were detected, consistent with the higher production rate of SVPs by these cells, whereas P-protein and core-specific peptide signals were absent. No or only background levels of HBV-specific peptides were identified in the IP conducted with the IgG control antibody or using samples derived from HepG2 cells.

Regarding host cell proteins co-purifying with HBV virions and fSVPs, a total of 790 host proteins was detected (Table S2A). Principal component analysis (PCA) of the individual samples confirmed high correlation within biological replicates (Fig. S4A). For further analysis, only proteins consistently identified in at least 3 biological replicates of at least one experimental group were retained (n=269 host proteins; Table S2B). This group of enriched host proteins was additionally filtered to retain only proteins completely missing from all the IgG control groups or displaying at least a median log2 (fold-change) ≥ 3 for each HBs-specific pull-down when compared to the corresponding IgG control group. In this way, 168 host cell proteins were identified, especially in the fSVP fractions derived from HepG2-HB2.7 cells, consistent with their higher SVP production rate as compared to HepAD38 cells (Fig. 1D and Table S2B). Although this higher number of input particles allowed the detection of even low amounts of host cell proteins in this fSVP fraction, a substantial subset of the proteins was also detected in virion and fSVP fractions from HepAD38 cells. Among all the identified host proteins, chaperones of different families were enriched in all conditions. Several of them have been previously implicated in HBV biology, including HSPB1 modulating cccDNA activity, and HSC70 binding to PreS1 and regulating L protein topology (Fig. S4B)^22, 23^. Three proteins, CASP14, CWF19L2 and HNRNPC were highly enriched in virion fractions with little to no signal overlap with peptides detected in the fSVP preparations (Fig. 1D and Table S2B). Moreover, members of the apolipoprotein family, such as ApoA1, ApoA2, ApoC1, ApoC3 and ApoE, were detected in both virion and fSVP fractions derived from both cell lines (Fig. 1D and Fig. S4B).

Taken together, we identified a set of previously published host proteins, along with several hitherto unknown host cell proteins selectively enriched with highly purified HBV virions and/or fSVPs.

### Functional validation of virion- and fSVP-associated host proteins upon HBV infection

To determine whether proteins identified via the proteome analysis of virions and fSVPs contribute to the HBV replication cycle, 48 candidate host proteins were cherry-picked and validated by RNAi-based screening in the context of HBV infection (Fig. 2A). Briefly, two days after reverse transfection of siRNAs, HepG2-NTCP A3 cells were infected with HBV, and the levels of secreted HBeAg, HBsAg, intracellular core protein (HBcAg) and viral RNAs were measured at day 5 post infection (Fig. 2B-E and Fig. S5A-B). In addition, cytotoxicity of a given knock-down was determined at seven days after siRNA transfection, identifying only one knock-down as cytotoxic (PCBP1; Fig. S5C). For each parameter, values were normalized to those obtained with the non-targeting siRNAs pool (siNT). Cells treated with a siRNA targeting viral transcripts (siHBV) or the peptidic entry inhibitor Bulevirtide (MyrB) were used as positive controls; both reduced HBV parameters without cytotoxic effects (Fig. 2B-E, Fig. S5). Phenotypes were ranked based on secreted HBeAg levels, a sensitive HBV replication marker. Overall, results of the different HBV replication parameters were consistent, although changes of HBsAg levels were less pronounced while intracellular core protein measurements had higher variability as compared to HBeAg. While depletion of several host proteins enhanced HBV infection, we focused on HBV host-dependency factors whose knockdown reduced viral parameters. In this regard, the most consistent and strongest effect was found with ApoC1 knockdown decreasing all viral parameters (protein and RNA) in a very reproducible manner (Fig. 2B-E).

**Fig. 2.**
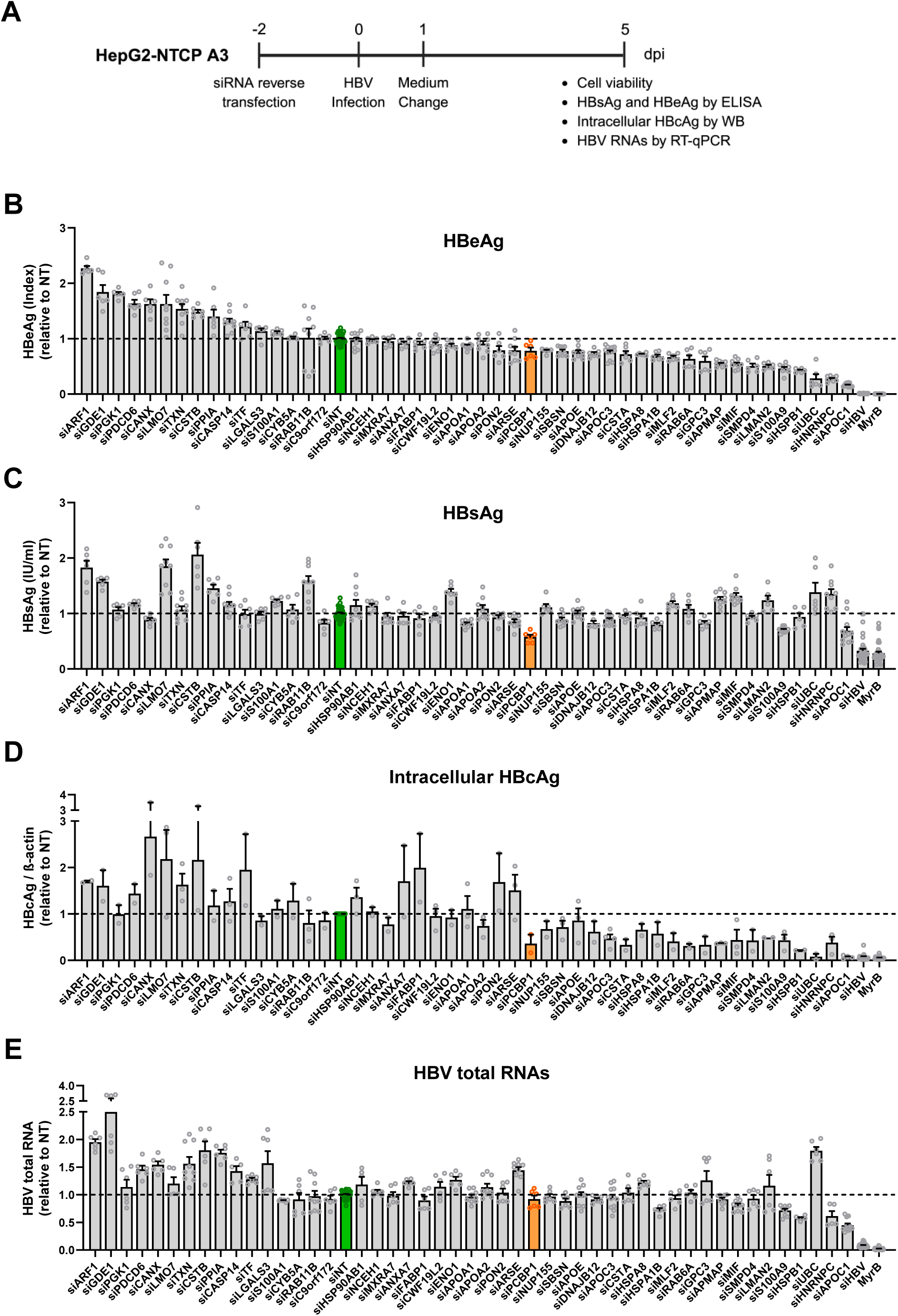
siRNA-based screen identifies novel HBV host-dependency factors. (A) Schematic of the experimental approach. HepG2-NTCP A3 cells were reverse-transfected with gene-specific siRNAs pool for 48 h and subsequently infected with HBV (500 genome equivalents (GE)/cell with 4% PEG8000). After 24 h, the inoculum was removed, cells were washed and cultured for an additional 5 days. Cells treated with the entry inhibitor Bulevirtide (MyrB) or transfected with siRNA targeting HBV transcripts (siHBV) served as positive controls. (B-C) HBV infection was evaluated by measuring secreted HBeAg and HBsAg at 5 days post infection (dpi) by ELISA. (D) Intracellular HBcAg was detected by western blot and quantified using ImageLab. (E) HBV total RNAs were quantified by RT-qPCR. Values were normalized to cells transfected with non-targeting siRNA (siNT), set to 1 (green bar). Data are presented as mean ± SEM from at least two independent experiments performed in triplicate. Orange bar, flawed result because of cytotoxicity (see Fig. S5C).

As noted above, five human apolipoproteins, ApoA1, ApoA2, ApoC1, ApoC3 and ApoE, were identified in our proteomic analysis of HBV virions and fSVPs derived from either cell line. Amongst these, ApoE was previously reported to enhance HBV infection and production^16^. Therefore, we compared the roles of the apolipoproteins identified in our proteome for their impact on HBV infection/replication and the production of SVPs, the latter from an integrated HBV DNA genome fragment (cell line HepG2-HB2.7). In HBV infection experiments using HepG2-NTCP A3 cells, knockdown efficiency measured seven days after siRNA transfection was high (Fig. 3A). Zooming into the RNAi validation results (Fig. 2), amongst the five apolipoproteins, ApoC1 knockdown caused the strongest reduction of HBV parameters, i.e. secreted HBsAg and HBeAg (Fig. 3B), intracellular HBV transcripts and pgRNA (Fig. 3C) as well as intracellular core protein (Fig. 3D), all determined five days after infection. In the context of the HB2.7 cell line, extracellular HBsAg was also reduced most by ApoC1 knockdown (Fig. 3E). These results suggest that ApoC1 plays a role not only in HBV infection/replication, but also, at least to some extent, in the production or release of SVPs from integrated viral sequences.

**Fig. 3.**
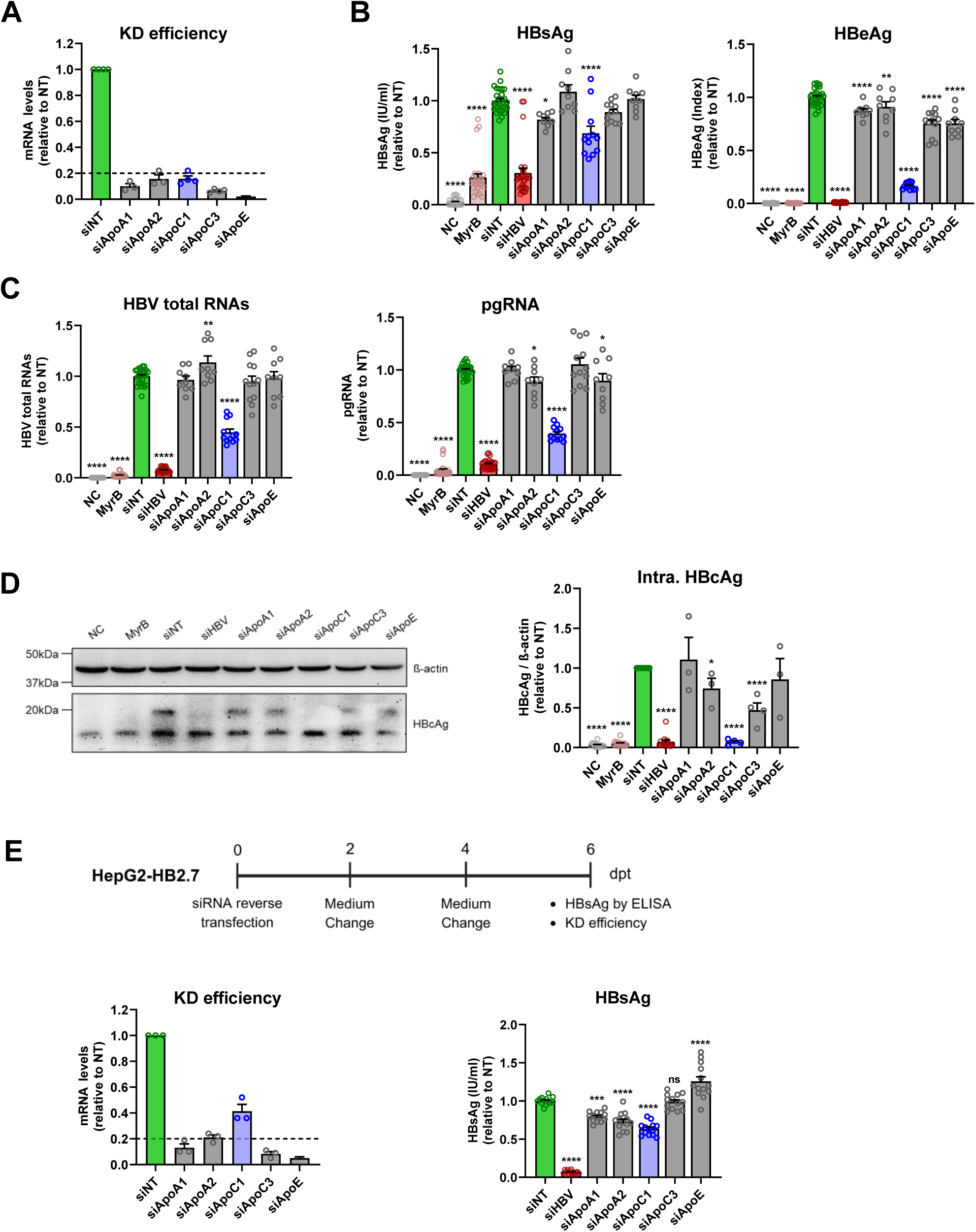
Comparative analysis of the impact of apolipoprotein depletion on HBV infection and HBsAg production. (A-D) Detailed view of apolipoprotein knockdown results from Fig. 2. (A) Knockdown efficiency was assessed by RT-qPCR. (B) Secreted HBsAg and HBeAg were measured by ELISA. (C) HBV total RNAs and pgRNA were quantified by RT-qPCR. (D) Intracellular HBcAg was detected by western blot and quantified using Image Lab. One out of at least three independent experiments is shown. (E) HepG2-HB2.7 cells were reverse-transfected with siRNAs; medium was refreshed after 48 h. Cells were harvested 6 days post transfection. Knockdown efficiency and secreted HBsAg were measured by RT-qPCR and ELISA, respectively. Values were normalized to siNT, set to 1 (green bar). Data represent mean ± SEM from at least three independent experiments. B, C, D, E, One-way ANOVA; *p<0.05, **p<0.01, ***p<0.001, ****p<0.0001, ns-not significant (p>0.05). NC, no infection control; MyrB, Bulevirtide-treated cells.

### ApoC1 is a proviral host protein promoting the HBV replication cycle

ApoC1 is the smallest, exchangeable, and most positively charged apolipoprotein. It regulates the activity of several lipid metabolizing enzymes and the function of lipoprotein receptors, including SR-B1^24^. Since ApoC1 was identified as top hit from the phenotypic screen, we further validated its relevance for the HBV replication cycle. HepG2-NTCP A3 cells were transfected with siRNAs targeting ApoC1, 48 h later cells were infected with HBV and the kinetics of secreted HBsAg, HBeAg, and HBV DNA were evaluated at day 5, 7 and 9 post infection. ApoC1 knockdown significantly reduced HBsAg and HBeAg, comparable to the NTCP knockdown control at all time points (Fig. 4A). In the case of extracellular HBV DNA, a marker for virus particle production, the signal detected at day 5 post infection was derived from input viruses, as levels were equivalent among all siRNA-transfected cells and cells treated with the entry inhibitor MyrB (Fig. 4A, lower panel). However, at later time points when most of the input had decayed, HBV DNA was significantly reduced upon ApoC1 knockdown relative to the siNT-transfected control cells, with the reduction being comparable or even stronger to what we observed upon NTCP depletion. Further quantification of the number of infected cells by immunofluorescence and cccDNA levels by qPCR confirmed the ApoC1 knockdown phenotype (Fig. 4B, C, respectively). The same result was found when we conducted HBV infection in the absence of polyethylene glycol (PEG) that is commonly used to enhance the binding and internalization of virions to cells (Fig. S6). Finally, we corroborated the role of ApoC1 in promoting HBV infection/replication in two independent ApoC1 knockout cell clones derived from HepG2-NTCP A3 cells (Fig. S7), thus making any off-target effects very unlikely.

**Fig. 4.**
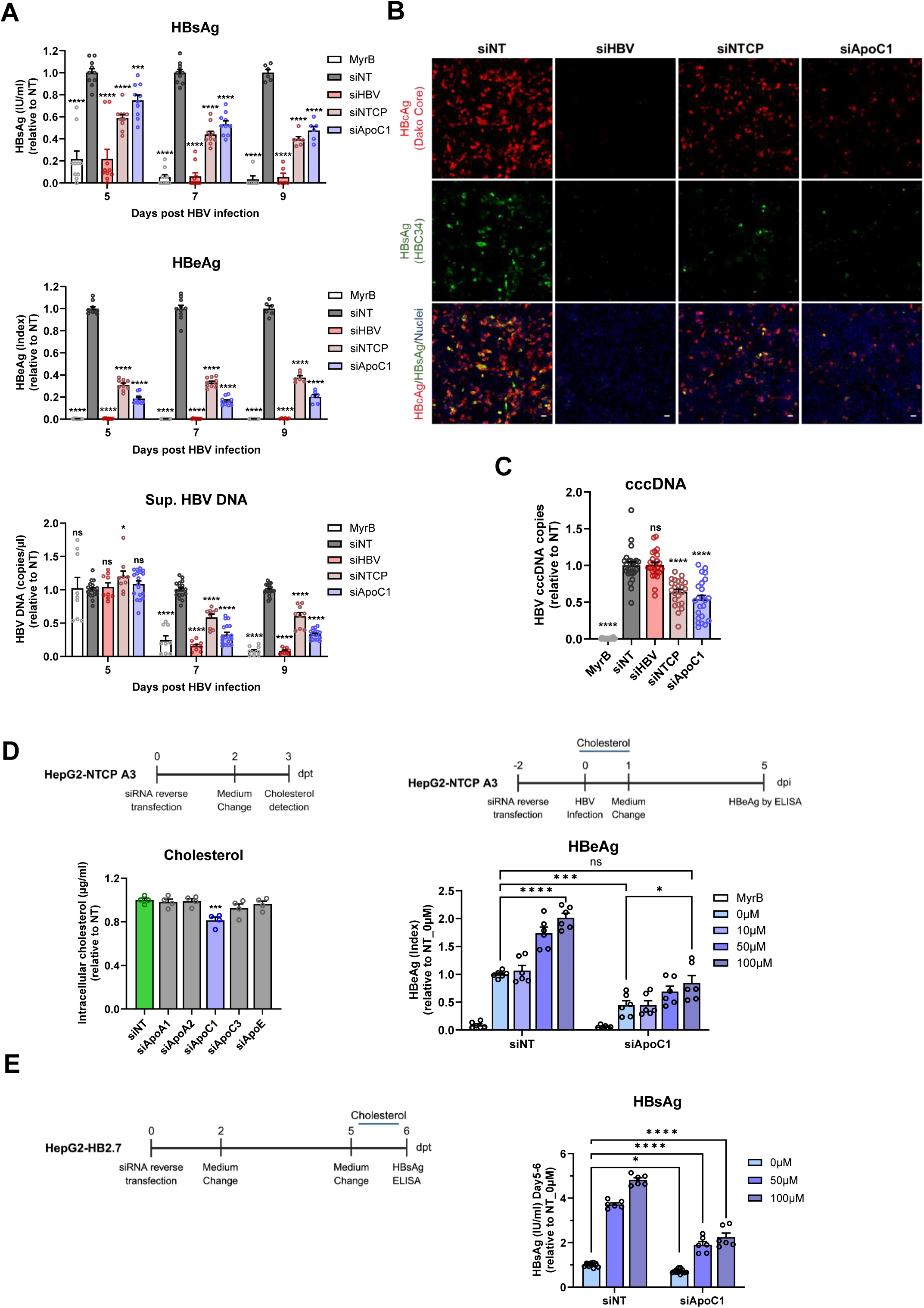
Role of ApoC1 for HBV infection and intracellular cholesterol level. (A-C) HepG2-NTCP A3 cells were reverse-transfected with siRNAs for 48 h and then infected with HBV (500 GE/cell; 4% PEG8000). After 24 h, the inoculum was removed, cells were washed and harvested at different time points. (A) Secreted HBsAg, HBeAg and HBV DNA were quantified by ELISA and qPCR at 5, 7 and 9 dpi, respectively. (B) Intracellular HBc and HBs proteins were assessed by immunofluorescence at 5 dpi. Red, green and blue signals depict HBc protein, HBs protein and nuclei, respectively. Scale bar: 100µm. (C) HBV cccDNA was detected by qPCR at 5 dpi. (D) HepG2-NTCP A3 cells were reverse-transfected with siRNAs, and medium was replaced by FBS-free medium after 48 h. Left: Intracellular cholesterol levels, measured one day later. Right: Cholesterol supplementation during HBV infection; secreted HBeAg was measured at 5 dpi. (E) HepG2-HB2.7 cells were reverse-transfected with siRNAs and 48 h later medium was refreshed. Cells were treated with water-soluble cholesterol between day 5 and 6 and secreted HBsAg was quantified. Data represent mean ± SEM from at least three independent experiments. A, D (right panel), E, Two-way ANOVA; C, D (left panel), One-way ANOVA; *p<0.05, ***p<0.001, ****p<0.0001, ns-not significant (p>0.05).

Since previous studies have reported functional redundancy of ApoC1 and ApoE^25^, we investigated whether ApoE depletion would have additional or even synergistic effect with ApoC1 depletion.

ApoC1 knockout cells were transfected with ApoE-specific siRNAs and subsequently infected with HBV (Fig. S8A). Secreted HBsAg and HBeAg, as well as intracellular HBc protein, remained comparable in ApoC1 knockout cells, regardless of additional ApoE knockdown (Fig. S8B, C). Similar results were observed in knockdown assays comparing ApoC1 single knockdown and ApoC1/ApoE double knockdown on HBV infection (Fig. S8D). These results suggested that ApoE could not compensate ApoC1’s role in promoting the HBV replication cycle.

### ApoC1 regulates intracellular cholesterol level affecting HBV and HDV entry

Cholesterol plays an important role in the HBV replication cycle^26^. Moreover, ApoC1 has been reported to regulate intracellular cholesterol levels^27^. Therefore, we quantified cholesterol amounts in HepG2-NTCP A3 cells depleted for ApoC1 or any of the other 4 different apolipoproteins identified in our proteome (Fig. 4D, left panel). Interestingly, we observed a small, but significant reduction of cholesterol only upon knockdown of ApoC1. To determine whether intracellular cholesterol amounts affect the HBV replication cycle, we conducted rescue experiments by replenishing ApoC1 knockdown cells with different concentrations of water-soluble cholesterol that was added to the cell culture medium. Although we observed an increase of the HBV replication marker HBeAg in ApoC1 knockdown cells by exogenously added cholesterol (around 2-fold at high cholesterol concentration), an analogous increase was found in cells transfected with siNT (Fig. 4D, right panel). This result indicated that cholesterol is a more generally limiting factor in the HBV replication cycle and not specific to ApoC1 depletion. Nevertheless, the reduction of cholesterol we observed in ApoC1 knockdown cells could explain, at least in part, the impairment of HBV replication observed in these cells. Indeed, cholesterol addition restored HBV replication close to the level detected in siNT-transfected control cells (Fig. 4D, right panel). Moreover, we found that cholesterol substitution also boosts SVP production in the context of HB2.7 cells (Fig. 4E) arguing that cholesterol supports several steps of the HBV replication cycle, including particle production.

### Association of ApoC1 with HBV virions and filamentous SVPs

In addition to regulating intracellular cholesterol levels, ApoC1 might promote the HBV replication cycle via association with virions and fSVPs, consistent with its detection in our proteome (Fig. 1D). To address this assumption, we analyzed two independent heparin-purified cell culture-derived HBV stocks by HBs- or ApoC1-specific IP and quantified the captured particles by Western blot and qPCR (Fig. 5A). We observed co-IP of ApoC1, HBc protein and HBV DNA when we captured the particles with an HBs-specific antibody; vice versa, co-IP of HBc protein and viral DNA was found when we captured the particles with the ApoC1-specific antibody (Fig. 5A). The analogous result was found using supernatants of Huh7 cells transfected with a plasmid containing a 1.1-overlength HBV genome, thus excluding confounders due to HepG2 cells or heparin purification (Fig. 5B and Fig. S9A). Notably, analysis of HBV particles contained in sera of three patients with CHB corroborated the association of ApoC1 with HBV particles (Fig. 5C and Fig. S9B).

**Fig. 5.**
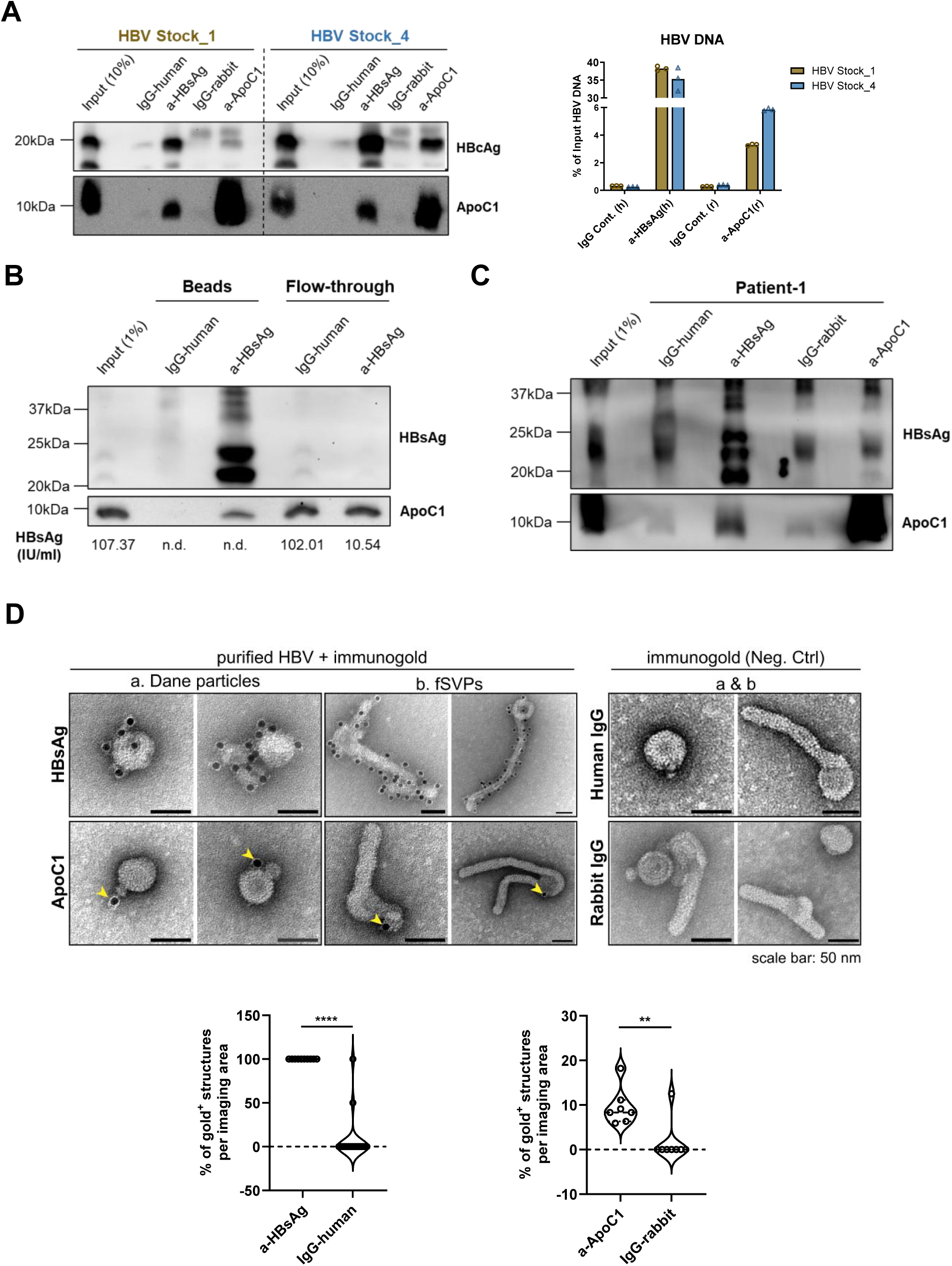
ApoC1 associates with HBV virions and filamentous SVPs. (A) Two different heparin-purified HBV stocks were subjected to immunoprecipitation using HBsAg- or ApoC1-specific or IgG antibodies. Eluates were analyzed for ApoC1 and HBcAg by western blot or HBV DNA by qPCR (normalized to input). (B) Huh7 cells were transfected with a plasmid containing a 1.1- overlength HBV genome. HBV accumulating in culture supernatant between day 4 and 7 post transfection were subjected to immunoprecipitation with HBsAg-specific or IgG antibodies. HBsAg ELISA values of input and flow-through fractions are indicated on the bottom. N.d., non-detectable. (C) HBV particles in serum of a patient with CHB were analyzed by immunoprecipitation and western blot. (D) HBV virions from HepAD38 cells purified by heparin affinity chromatography, size exclusion chromatography and sucrose gradient centrifugation were immunogold-labeled for HBsAg and ApoC1 and visualized by transmission electron microscopy. IgG antibodies served as specificity controls. Quantification of gold-positive structures is shown below. Scale bar: 50 nm. Unpaired t test; **p<0.01, ****p<0.0001.

To further corroborate the association of ApoC1 with HBV particles, HepAD38 culture-derived and purified HBV virions or fSVPs were labeled with ApoC1- or HBs-specific primary antibodies, followed by Protein A-coupled to 10 nm gold particles and samples were analyzed by transmission electron microscopy (TEM). As expected, HBs proteins were present on the surface of virions and fSVPs (Fig. 5D and Fig. S9C). Notably, ApoC1 was also detected in association with highly purified virions and fSVPs. Quantification of gold-labeled particles revealed ∼10% of the HBV particles being ApoC1 positive (Fig. 5D). Interestingly, HDL-like structures were occasionally observed adjacent to HBV virions and fSVPs, indicating an association of HBV particles with ApoC1-containing lipoprotein particles (Fig. 5C and Fig. S9C).

Taken together, these results confirm ApoC1 association with HBV virions and fSVPs. Our data indicate that at least a fraction of ApoC1 associates with these particles in the form of HDL.

### ApoC1 regulates HBV and HDV entry

To study the functional role of ApoC1 in HBV uptake into liver cells, we performed ApoC1 antibody-mediated neutralization assays in the context of HBV infection. Strikingly, cells infected with purified HBV stocks that had been pre-incubated with an ApoC1-specific antibody released a dose-dependently lower amount of HBsAg, HBeAg and HBV DNA into the culture supernatant, correlating with reduced amounts of intracellular HBc protein (Fig. 6A). This result suggested that viral entry is facilitated by HBV-associated ApoC1.

**Fig. 6.**
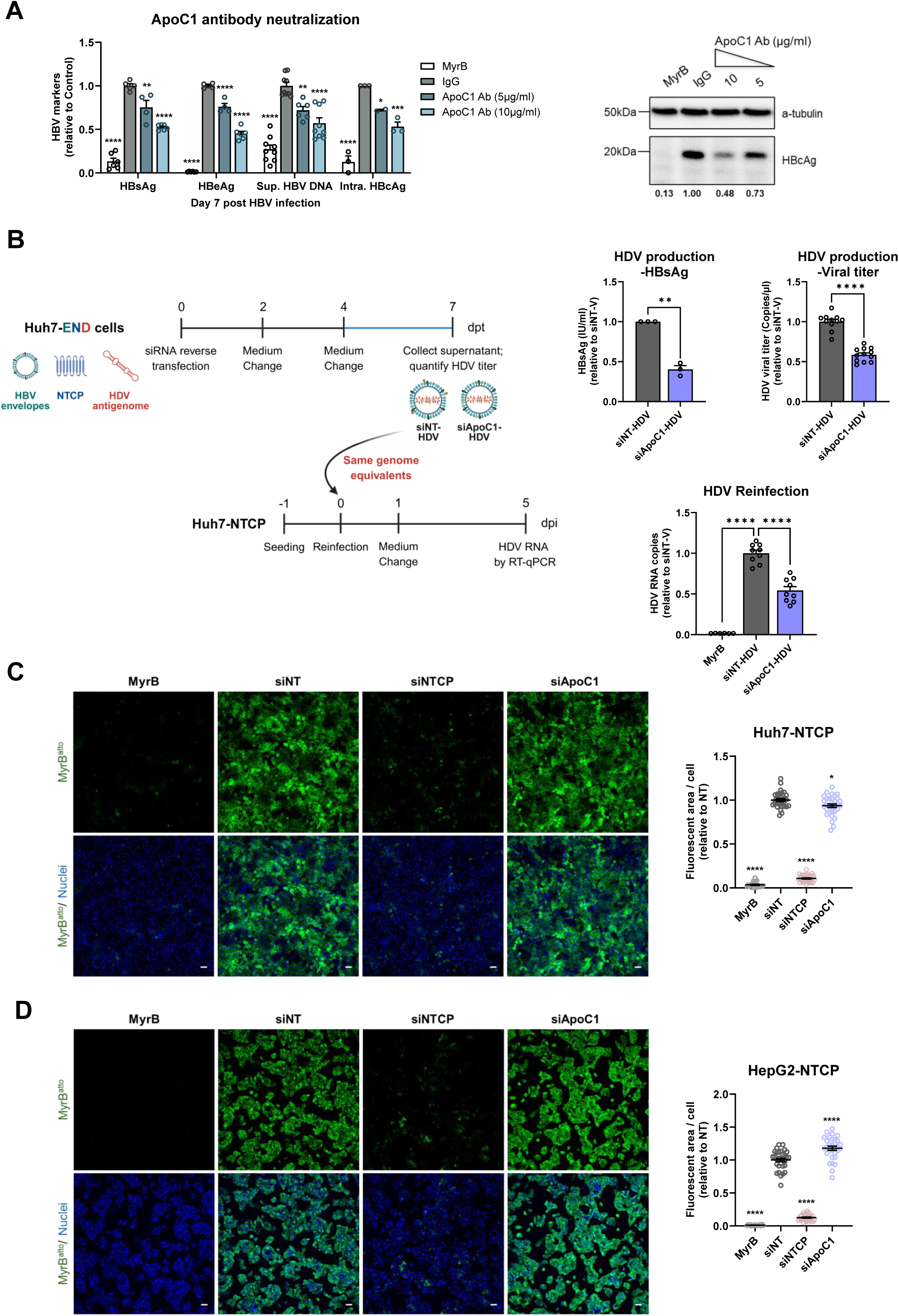
ApoC1 enhances HBV and HDV infectivity. (A) HBV stocks were pre-incubated with ApoC1 antibody for 1 h at room temperature prior to infecting HepG2-NTCP A3 cells (1,500 GE/cell) in the absence of PEG8000 and FBS for 24 h. Secreted HBsAg, HBeAg and HBV DNA were quantified by ELISA and qPCR at 7 dpi, respectively. Intracellular HBcAg (right panel) was detected by western blot at 7 dpi and quantified using Image Lab. One out of three representative experiments is shown. (B) To obtain HDV depleted for ApoC1, Huh7-END cells (schematic on the left) were transfected with siApoC1 or siNT. HBsAg levels and HDV titers were determined by ELISA and RT-qPCR, respectively. Owing to lower HDV production in ApoC1 depleted cells (upper right panels), Huh7-NTCP target cells were inoculated with the same HDV genome equivalents to evaluate the impact of ApoC1 depletion on HDV infectivity. At 5 dpi, intracellular HDV RNAs were quantified by RT-qPCR (lower right panel). Data represent mean ± SEM from three independent experiments. (C) Huh7-NTCP or (D) HepG2-NTCP A3 cells were reverse-transfected with given siRNAs for 48 h and subsequently exposed to 200 nM atto488-labeled MyrB peptide (MyrB^atto^) for 30 mins at 37°C. Green and blue signals on the left panels depict MyrB^atto^ and nuclei, respectively. Cells incubated with an excess of unconjugated MyrB peptide served as nonspecific binding control. Scale bar: 100µm. Data represent mean ± SEM from two independent experiments performed in duplicate. A, B (lower right panel), C, D, One-way ANOVA; B (upper panels), unpaired t test; *p<0.05, **p<0.01, ***p<0.001, ****p<0.0001, ns-not significant (p>0.05).

HBV and its satellite virus HDV share the same envelope and thus, the early steps of hepatocytes infection. Consistently, we found that ApoC1 knockdown impaired HDV infection (Fig. S10). Taking advantage from this observation, we used HDV as a tractable surrogate to determine how ApoC1 depletion affects virion infectivity. We employed Huh7-END cells^28^ stably releasing HDV and depleted for ApoC1 by using siRNA-mediated knockdown (Fig. 6B). HDV released from these cells and from Huh7-END cells transfected with siNT was quantified for viral genomes and equal amounts of genome equivalents were used to infect permissive Huh7-NTCP cells. We found that ApoC1-deficient HDV had significantly lower specific infectivity as compared to wildtype HDV virions (Fig. 6B) arguing that ApoC1 associated with HDV, and -by analogy- with HBV, increases infectivity of the virus particles.

Both HBV and HDV specifically bind to NTCP, followed by internalization into hepatocytes via receptor-mediated endocytosis. To assess whether, in addition to increasing virion infectivity, ApoC1 knockdown affects cell surface availability of NTCP, we employed a fluorophore-conjugated preS1-peptide^4^. Using HepG2 or Huh7 cells stably expressing NTCP, ApoC1 depletion had no effect on NTCP cell surface abundance (Fig. 6C, D).

### Impairment of HBV infection by pharmacological inhibition of SR-B1

Since ApoC1 is mainly associated with HDL (Fig. 7A), the role of HDL in HBV infection was investigated. Pretreatment of purified HBV stocks with exogenous HDL increased infection as deduced from elevated amounts of HBsAg and HBeAg secreted from infected cells, indicating enhanced viral infectivity by HDL (Fig. 7B).

**Fig. 7.**
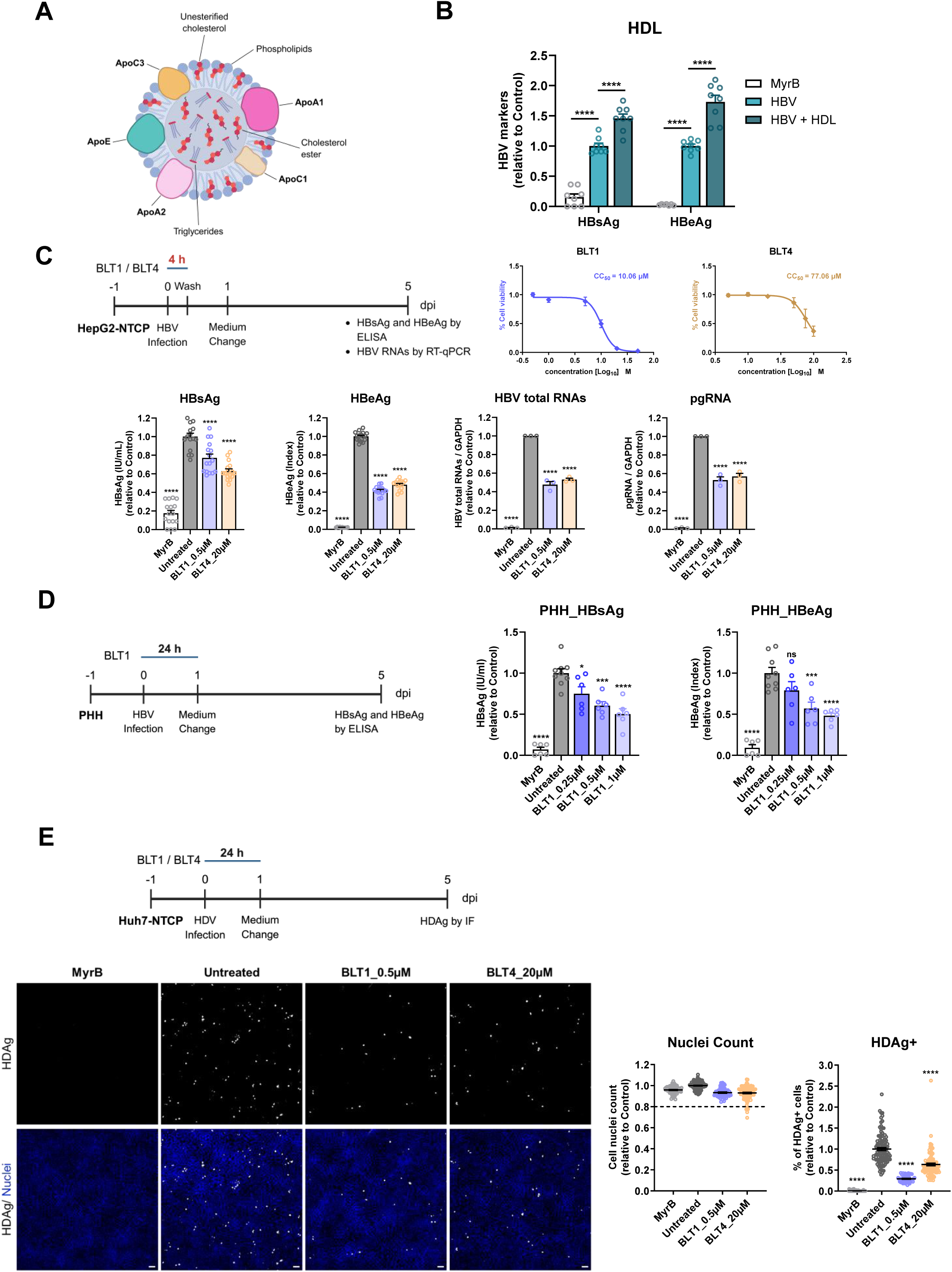
Inhibition of HBV and HDV infection by pharmacological block of SR-B1. (A) Schematic diagram of high-density lipoprotein (HDL). Created in BioRender. (B) Heparin purified HBV stocks were pre-incubated with HDL (300 µg/ml) for 30 mins at RT prior to infecting HepG2-NTCP A3 cells (1,500 GE/cell) in the absence of 4% PEG8000. After 24 h in FBS-free medium, cells were washed and complete medium was added. Secreted HBsAg and HBeAg were quantified at 7 dpi. (C) HepG2-NTCP A3 cells or (D) PHHs were treated with SR-B1 inhibitors (BLT1 or BLT4) throughout infection with HBV (1,000 GE/cell with 4% PEG8000). Secreted HBsAg, HBeAg, HBV total RNAs and pgRNA were quantified by ELISA and RT-qPCR at 5 dpi, respectively. (E) Huh7-NTCP cells were treated with BLT1 or BLT4, starting at 4 h prior to and throughout infection with HDV (5 GE/cell with 4% PEG8000). After 24 h, cells were washed and cultured in medium without SR-B1 inhibitors. Intracellular HDAg was assessed by immunofluorescence at 5 dpi. White and blue signals depict HDAg and nuclei, respectively. Quantification of nuclei number and HDAg-positive cells are shown. Scale bar: 100µm. Data are presented as mean ± SEM from three independent experiments. B, C, D, E, One-way ANOVA; *p<0.05, ***p<0.001, ****p<0.0001, ns-not significant (p>0.05).

Since SR-B1 is a natural receptor of HDL, we next studied the possible contribution of SR-B1 to HBV infection. Earlier studies reported that HBV infection is facilitated by ApoE, which binds to LDL receptors^29^. We therefore compared the impact of SR-B1 depletion on HBV infection with the one of LDL and VLDL depletion. As shown in Fig. S11A, knockdown of SR-B1 significantly reduced HBV infection. A comparable effect was found for LDL receptor depletion whereas knockdown of the VLDL receptor had no effect. Combination of SR-B1 and LDL receptor knockdown had no additive effect. Analogous results were obtained upon silencing of these receptors in HDV infection, consistent with the same entry route used by both viruses (Fig. S11B).

SR-B1 mediates bidirectional lipid transport between HDL and cells. One of its primary natural functions is to facilitate the selective uptake of cholesterol esters from HDL into liver cells for bile acid synthesis and the efflux of the free cholesterol to HDL^30, 31^. We explored this function of SR-B1 in HBV infection using the known pharmacological inhibitors BLT (block lipid transport)-1 and BLT-4. These small molecules bind to SR-B1 in an irreversible or reversible manner, respectively, and inhibit SR-B1 mediated lipid transfer activity between HDL and cells ^32, 33^. HepG2-NTCP A3 cells were infected with HBV for 4 h in the presence of BLT compounds and viral replication was measured 5 days later. Both BLT compounds significantly reduced HBV replication, indicating that SR-B1 is a potential co-factor for HBV entry (Fig. 7C). To corroborate this conclusion in a more relevant cell system, the effect of BLT-1, the more potent and specific inhibitor^32, 34^, was evaluated in HBV-infected primary human hepatocytes (PHHs) derived from two different donors (Fig. 7D). The dose-dependent reduction of extracellular HBsAg and HBeAg supported our conclusion on the important role of SR-B1 in HBV entry. Consistently, HDV infection was also reduced by both SR-B1 inhibitors (Fig. 7E).

Taken together, these results suggest that HDL enhanced HBV and HDV infection in an ApoC1-and SR-B1-dependent manner. ApoC1, probably via HDL, targets virions to SR-B1 that might act as co-factor to facilitate successful infection. This would explain why supplementation of exogenous HDL enhanced HBV infection while pharmacological inhibition of SR-B1 suppressed the infection.

## Discussion

By using a sequential purification approach and quantitative proteomics, we comprehensively determined the first proteome composition of HBV particles allowing the detection of very low abundance proteins such as the HBV P-protein that is present at approximately one copy per DNA-containing virion^21^. Consistent with earlier studies, the HBx protein was not detected^35^. Regarding host cell proteins, a recent cryo-EM study of pgRNA-containing nucleocapsids revealed an electron-dense region within the capsid interior. In addition to the pgRNA and the HBV polymerase, this region was proposed to contain entrapped host factors^36^. Consistently, we identified three cellular proteins (CASP14, CWF19L2 and HNRNPC) that were highly enriched in the virion/fSVP preparation derived from HepAD38 cells, but not in fSVPs from HepG2-HB2.7 cells. However, whether these proteins play a specific role in the HBV replication cycle, but not in SVP production remains to be determined. We also detected several chaperones in the virion and fSVP fractions. Amongst them is HSP90AB1, a component of the HSP90 complex, earlier shown to facilitate HBV polymerase-epsilon RNA interactions and assembly of the reverse transcription initiation complex^37^.

Additional chaperones that we identified in association with the viral envelope proteins, included HSC70 and its interaction partner, the cytosolic anchorage determinant binding protein. These proteins were earlier shown to inhibit co-translational translocation of the PreS1 domain through the viral envelope enabling the dual topology of the L protein^23^. Similarly, PPIA has been reported to interact with HBsAg and is upregulated in the sera of patients with CHB^17^. In our fSVP proteome, we also detected the HSP40 chaperone DNAJB12, a reported interaction partner of the nucleic acid polymer REP-2139 that interferes with the assembly and release of HBV SVPs and is currently in clinical trials for HBV and HDV infections^38^. These findings underscore the superior properties of our particle purification method.

One intriguing feature of our proteome is the presence of five HDL-enriched apolipoproteins co-purifying with fSVPs. HDL mediates the transport of cholesterol from peripheral tissues to the liver by carrying both free cholesterol and cholesteryl esters as part of the reverse cholesterol transport pathway. Recently, macrophages were shown to internalize lipoprotein-associated HBV particles into recycling endosomes enriched in cholesterol, which are subsequently trafficked back to the cell surface to facilitate hepatocyte infection^18^. The accumulation of HDL apolipoproteins together with cholesterol in both the HBV lipid envelope and host endosomal membranes may therefore confer favorable biophysical properties promoting viral entry. Cholesterol influences membrane fusion in several enveloped viruses, including Ebola virus, HIV, Zika virus and SARS-CoV-2^39–42^. Consistently, cholesterol-depleted HBV virions treated with methyl-β-cyclodextrin have reduced virion diameter and lower infectivity, both of which were restored upon cholesterol replenishment^26^. In our study, HDL supplementation during infection enhanced viral entry, further supporting a role for lipoproteins during the early stages of HBV infection (Fig. 7B). Moreover, pharmacological inhibition of cholesterol and cholesteryl ester transport through the HDL receptor SR-B1 reduced HBV and HDV infection (Fig. 7C-E). While these results suggest that SR-B1 might act as an auxiliary receptor of HBV, consistent with an earlier study^43^, we note that SR-B1 splice variants have been reported to localize to distinct subcellular compartments, including the plasma membrane and endosomes^44^. This localization may extend the role of SR-B1 beyond its interaction with HBV particles at the cell surface. Moreover, SR-B1 association with lipoproteins is not limited to HDL as interactions with LDL and VLDL have also been reported^44, 45^. Therefore, we cannot exclude the involvement of other lipoproteins in HBV entry, which might explain the earlier report of LDLR being an important host cell factor for HBV infection^29^.

In our comparative analysis of the 5 different apolipoproteins identified in our proteome, ApoC1 depletion had the strongest negative effect on HBV. The impact was even stronger than depletion of ApoE previously reported to enhance HBV attachment to the cell surface by binding to the HSPG syndecan-2 and the LDLR^29, 46^. By using ApoC1-depleted HDV as surrogate for HBV, we observed lower infectivity of the virus particles suggesting that virion associated ApoC1 enhances entry (Fig. 6B). Interestingly, a role for ApoC1 in viral membrane fusion has been demonstrated previously for HCV^47^ and a similar mechanism might operate during HBV infection. However, owing to the lack of a robust HBV fusion assay and the limited efficiency of current in vitro infection systems, this hypothesis cannot be addressed experimentally.

The bovine genome lacks an ApoC1 gene^48^, excluding the possibility of ApoC1 exchange between fetal bovine serum and HBsAg in cell culture, although exchange with lipoproteins released from cultured cells cannot be excluded. We also detected association between ApoC1 and HBV virions or SVPs in sera from patients with CHB, indicating that HBV can hijack circulating lipoproteins possibly to facilitate its transport to the liver. Given these findings, combining current HBV therapies with FDA-approved serum lipid-modulating drugs may represent a novel antiviral strategy.

## Data Availability

All data supporting the findings of this study are available within this paper and its supplementary information. The mass spectrometry proteomics data have been deposited to the ProteomeXchange Consortium via the PRIDE^49^ partner repository with the dataset identifier PXD076462. Source data are provided with this paper (Table S1 and S2).

## Acknowledgements

We acknowledge the excellent support with proteomics measurements by Dr. Antonio Piras (Institute of Virology, Technical University of Munich). We are grateful to Uta Haselmann for EM sample preparations. We gratefully acknowledge the Electron Microscopy Core Facility (EMCF) of Heidelberg University, headed by Charlotta Funaya, for expert support and access to their equipment. HDL illustration in Fig. 7A was created in BioRender https://BioRender.com/bzu8lac.

## Abbreviations

## Author contributions

Conceived and designed the experiments: SY, FN, SS, RB. Performed the experiments: SY, FN, MP, PS. Analyzed and interpreted the data: SY, FN, PS, RB. Contributed reagents/materials/analysis tools: YL, AP, PS, SN, SU, SS, RB. Drafted the manuscript: SY, FN, RB.

