## Supplementary Figures and Tables for "Hepatitis B virus proteome analysis identifies apolipoprotein C1 facilitating particle production and virus entry": Supplementary Materials-2026.04.02_Final.pdf

### Contributed equally to this work

Fig. S6 ........ 11
Fig. S7 ........ 11
Fig. S8 ........ 18
Fig. S9 ........ 19
Fig. S10 ........ 21
Fig. S11 ........ 22
Table S1 (Viral Peptides) ........ 24
Table S2 (Host Proteins) ........ 24
Table S3 (Primers) ........ 25
Table S4 (Antibodies) ........ 26
References ........ 28

#### Supplementary materials and methods

##### *Cell culture*

Huh7, HepG2-NTCP A3, HepG2-HB2.7 and HEK293T cells were maintained in DMEM (Life Technologies, USA) supplemented with 10% fetal bovine serum (FBS; Capricorn Scientific, Germany), 1x non-essential amino acids (Life Technologies, USA), 100 units/ml penicillin, 100 µg/ml streptomycin (Gibco, USA), and incubated at 37°C with 5% $CO_2$ and 95% humidity. HepAD38 cells were cultured in DMEM/F12 (Gibco, USA) supplemented with 10% FBS, 1x non-essential amino acids, 100 units/ml penicillin, 100 µg/ml streptomycin, 2 mM L-glutamine (Gibco, USA), 1 mM sodium pyruvate (Gibco, USA), 5 µg/ml insulin (SAFC Biosciences, UK) and 50 µM hydrocortisone (Sigma-Aldrich, USA). To suppress HBV production, 1 µg/ml doxycycline (Sigma-Aldrich, USA) was added to the HepAD38 medium for general maintenance of the cells.

Long-term cultures of HepG2, HepG2-HB2.7 and HepAD38 cells were performed in 875 $cm^2$ surface cell culture multi-flasks (Corning, USA) using DMEM/F12 containing 10% FBS or, for the fourth and fifth proteome sample production, 5% FBS. The cultured supernatants were collected every four days, sterile filtered through 0.45 µm membranes and stored at 4°C until further processing.

Cryopreserved primary human hepatocytes (PHHs) were purchased from Thermo Fisher Scientific and cultured according to the manufacturer's instructions.

##### *HBV particles purification*

HBV particles were purified using the Äkta purifier 100 and Äkta pure chromatography systems (Cytiva, UK) coupled to density gradient centrifugation and HBs-specific immunoprecipitation. In brief, HBV particles contained in filtered culture supernatants were loaded onto a HiTrap 5 ml heparin HP affinity column (Cytiva, UK), washed with TN buffer (20 mM Tris-HCl, 140 mM NaCl, pH 7.4) and eluted with a linear gradient of 140 mM to 2140 mM NaCl solution. After elution, the three peak fractions were pooled and further processed using a Superose 6 colunm (S6; Cytiva, UK) with a superloop. Subsequently, peak fractions eluted in the void volume, containing predominantly HBV virions and filamentous SVPs but lacking the majority of spherical SVPs, were subjected to a second round of heparin affinity chromatography using a HiTrap 1 ml heparin HP affinity column (Cytiva, UK). The concentrated peak fractions obtained from this column were layered on top of a discontinuous (60%, 45%, 35%, 15%) sucrose gradient and centrifuged at 164,000 × g overnight at 4°C using a SW60 rotor (Beckman Coulter, USA). Gradient fractions were collected from the bottom of the tube and the density and HBV protein content in each fraction was determined. Selected fractions were diluted in TN buffer and applied to Amicon Ultra 0.5ml centrifugal filters (Sigma-Aldrich, USA) to remove sucrose. The sucrose-free samples were subjected to immunoprecipitation using HBsAg- or IgG-specific antibodies non-covalently captured on magnetic beads. After three 15-minute washes with cold TN buffer on a rotator, followed by a brief wash with 100 mM Tris-HCl (pH 8.0) to remove residual salt, the beads were stored at −80 °C until use for proteomic analysis.

***Proteomics sample preparation, data analysis and processing***

Affinity-purified complexes on magnetic beads were denatured by incubation in 40 µl U/T buffer (6 M Urea, 2 M Thiourea, 100 mM Tris-HCl, pH 8.5), and reduction and alkylation carried out with 10 mM DTT and 55 mM iodoacetamide in 50 mM ABC buffer (50 mM $NH_4HCO_3$ in water, pH 8.0), respectively. After digestion with LysC (WAKO Chemicals, USA) (0.5 µg/sample) at room temperature for 3 h, the suspension was diluted in ABC buffer, and the protein solution was digested with 1 µg trypsin (Promega, USA) (IPs: 0.5 µg; whole-cell lysates: 1µg) overnight at room temperature. Digestion was stopped with 1% TFA, and peptides were purified on StageTips containing three C18 Empore filter discs (3M) and analyzed by liquid chromatography coupled to tandem mass spectrometry as previously described[1]. Briefly, purified peptides were loaded onto a 50-cm reverse-phase analytical column (75-µm diameter; ReproSil-Pur C18-AQ 1.Mµm resin; Dr. Maisch, Germany) and separated using an EASY-nLC 1200 system (Thermo Fisher Scientific, USA) with a 120 min gradient (80% acetonitrile, 0.1% formic acid; 5% (80% acetonitrile) to 30% for 95 min, 30% to 60% for 5 min, 60% to 95% for 5 min, wash out at 95% for 5 min, readjustment to 5% in 10 min) at a flow rate of 300 nl per min. Eluting peptides were directly analyzed on a Q-Exactive HF mass spectrometer (Thermo Fisher Scientific, USA) operated in data-dependent acquisition, including repeating cycles of one MS1 full scan (200-2,000 m/z, R=60,000 at 200m/z) followed by 15 MS2 scans of the highest abundant isolated and higher-energy collisional dissociation fragmented peptide precursors.

Raw MS data were processed with the MaxQuant software v.1.6.14 using the built-in Andromeda search engine to search against the human proteome (UniprotKB UP0000005640_9006, release

2017_10) containing forward and reverse sequences concatenated with the individual HBV open reading frames manually annotated, and searched with the label-free quantitation (LFQ) algorithm. Additionally, the intensity-based absolute quantification (iBAQ) algorithm and match between runs option were used. In MaxQuant, carbamidomethylation was set as fixed and methionine oxidation and N-acetylation and phosphorylation (pSTY) as variable modifications. Search peptide tolerance was set at 70 p.p.m. and the main search was set at 30 p.p.m. (other settings left as default). Search results were filtered with a false discovery rate of 0.01 for peptide and protein identification. Perseus software v.1.6.2.0 was used to process the data further. Protein tables were filtered to eliminate the identifications from the reverse database and common contaminants. When analyzing the MS data, only proteins identified on the basis of at least one peptide and a minimum of three quantitation events in at least one experimental group were considered. The iBAQ protein intensity values were log-transformed, normalized against the median intensity of each sample and shifted into positive numerical space by 12xlog2 values. Principal component analysis was used to identify and remove outliers. In total, three individual samples were removed from further downstream analysis and hit selection (HepAD38_HBV_anti-HBs_5, HepAD38_HBV_anti-IgG_4, HepAD38_SVP_anti-HBs_4).

To identify and remove outliers in principal component analysis, missing values were filled by imputation with random numbers drawn from a normal distribution calculated for each sample (width 0.3, down shift 1.8). For identification of high-confidence host proteins enriched in filamentous subviral particles (fSVP) or HBV virions (HBV), the pre-filtered matrix (n=269 host proteins) was further processed without any imputation step. The matrix was additionally filtered to retain only proteins completely missing from the IgG control group or displaying at least a median log2 (fold-change) $\geq 3$ for each pull-down when compared to the corresponding IgG control group (n=168 host proteins). To assess and visualize relative amounts of viral peptides across samples, absolute spectral intensities were extracted from the *peptides.txt* output table, log2-transformed and plotted without any further normalization.

##### *RNA and DNA transfection*

siRNAs for validation experiments were purchased from Dharmacon (USA) and reverse transfected using Lipofectamine RNAiMAX Transfection Reagent (Invitrogen, USA) according to the manufacturer’s instructions. Plasmid DNA transfection was performed using TransIT LT-1

(Mirus Bio, USA) according to the instructions of the manufacturer.

##### *HBV infection and quantification of viral antigens*

The HBV stock used for infection assays was prepared from cell supernatant of HepAD38 cells[2]. In brief, supernatants were collected and HBV particles (virions and SVPs contained therein) were purified by heparin affinity chromatography after removal of doxycycline. The virus titer was quantified by qPCR, and aliquots were stored at -80°C until use.

For infection, one day after seeding, HepG2-NTCP A3 cells were infected with HBV as indicated in each experiment. After washing with PBS, the cells were maintained in medium containing 2.5% DMSO (VWR International, USA). For Bulevirtide (MyrB) treatment, cells were pre-treated with the MyrB peptide (corresponding to amino acid residues 2-48 of the HBV preS1 domain) at 37°C for 30 mins and during infection.

Secreted HBsAg and HBeAg were determined by using the Architect assay (Abbott, Germany) and the ADVIA Centaur XP Immunoassay System (Siemens, Germany), respectively.

##### *HDV infection*

The HDV stock used for infection assay was prepared from cell culture supernatant of Huh7-END cells and virions and SVPs contained therein were purified by heparin affinity chromatography[3]. The virus titer was quantified by RT-qPCR, and aliquots were stored at -80°C until use.

For infection, one day after seeding, Huh7-NTCP cells were infected with 5 HDV genome equivalents per cell (MOI 5) in the presence of 4% polyethylene glycol (PEG) 8000 (Sigma-Aldrich, USA) and 1.5% DMSO for 24 h. After washing with PBS, the cells were maintained in medium containing DMSO for an additional 5 days.

##### *Cell viability assay*

Cell viability was evaluated using the WST-1 reagent (Roche, Switzerland) according to the manufacturer's instructions. The cells were incubated with freshly prepared WST-1 solution at 37°C for 30 mins and the absorbance at 450/620 nm was measured using an ELISA reader.

##### *DNA and RNA extraction*

Extracellular HBV DNA was extracted using the DirectPCR Lysis Reagent Cell (VWR International, USA) whereas HDV RNA was extracted from virus particles contained in the culture supernatant using the QIAamp Viral RNA Kit (QIAGEN, Germany). Total DNA and RNA from infected cells were isolated using the NucleoSpin Tissue kit (Macherey-Nagel, Germany) and the Monarch Total RNA Miniprep Kit (NEB, USA), respectively, according to the manufacturer's instructions. For HBV cccDNA preparation, total cellular DNA was treated with T5 exonuclease to remove all non-circular DNAs[4].

***PCR methods***

Extracellular HBV DNA amounts were determined using the iTaq universal SYBR Green Supermix (Bio-Rad, USA), while cccDNA quantification was performed using the PerfeCTa qPCR Toughmix (QuantaBio, USA) and values were normalized to ß-globin amplified in parallel. Total RNA was reverse transcribed using High-Capacity cDNA Reverse Transcription kits (Applied Biosystems, USA) and cDNAs were quantified and normalized to GAPDH analyzed in parallel. For absolute quantification, a 10-fold serial dilution of an HBV genome-containing plasmid or HDV genome-containing plasmid was used as standards. All qPCR experiments were conducted using a CFX96 thermocycler (Bio-Rad, USA).
The gene specific primers and probes used in the study are listed in Table S3.

***Western blotting***

Cells were lysed using 1x Laemmli buffer (6x Laemmli buffer stock: 97.5 mM Tris-HCl (pH 6.8), 30% glycerol, 3% SDS, 7.5% β-mercaptoethanol, 0.06% (w/v) bromophenol blue) and boiled at 95°C for 5-10 mins. Equal amounts of total proteins (cell lysates) or equal volumes of secreted proteins were separated by SDS-PAGE using a Mini-PROTEAN Tetra Cell (Bio-Rad, USA) and transferred to PVDF membranes (Bio-Rad, USA) using Trans-Blot Turbo transfer system (Bio-Rad, USA). The membranes were blocked with 5% BSA or 5% skim milk in PBS containing 0.1% Tween 20 for 1 h at room temperature, followed by incubation with primary antibodies overnight at 4°C. On the following day, the membranes were washed and incubated with secondary antibodies for 1 h at room temperature. After washing, protein bands were visualized using either the Clarity Western ECL substrate (Bio-Rad, USA) or Amersham ECL Prime Western Blot Detection Reagent (Cytiva, USA) on the Intas Chemocam system (Intas Science Imaging, Germany). For

fluorescence-labeled antibodies, the membranes were imaged using a LI-COR Odyssey CLx (LI-COR Biosciences, Germany) or an Intas Advanced Fluorescence Imager (Intas Science Imaging Instruments, Germany). Image J software was used for image processing, and band intensities were quantified using the Image Lab software package (Bio-Rad, USA).
The antibodies used in this study are listed in Table S4.

***Immunofluorescence microscopy***

HBV- and HDV-infected cells grown on coverslips were fixed with 4% paraformaldehyde (PFA) for 10-15 mins at room temperature. The cells were then permeabilized with 0.5% Triton X-100 for 10 mins and blocked with 3% BSA (Sigma-Aldrich, USA) in PBS for 1 h at room temperature. After blocking, cells were incubated with primary antibodies diluted in 0.5% BSA overnight at 4°C. On the next day, cells were washed with PBS and incubated with secondary antibodies together with DAPI (MoBiTec, Germany) for 1 h in the dark at room temperature, followed by mounting with Fluoromount-G mounting medium (Invitrogen, USA). Images were captured using a Nikon Eclipse Ti Inverted Microscope (Nikon Instruments Europe operations, Netherlands) and the NIS-Elements Advanced Research software and processed using the Image J software package.
The antibodies used in this study are listed in Table S4.

***Lentivirus production and generation of stable cell lines***

Lentivirus production was performed in HEK293T cells using the Calcium phosphate transfection kit (Takara, Japan). In brief, HEK293T cells were seeded in 10 cm diameter dishes one day before transfection. The pMD2.G plasmid encoding VSV-G glycoprotein (kindly provided by Prof. Didier Trono, EPFL, Switzerland) and lentiviral packaging plasmid pCMV-dR8.91 were co-transfected with pWPI expression vectors (for gene expression) or LentiCRISPRv2 vectors (Addgene #52961) harboring the guide RNA (for gene inactivation) at a 1:3:3 ratio. The medium was refreshed 24 h post transfection, and lentivirus containing supernatants were collected at 48 h, 72 h and 96 h post transfection, filtered through 0.45 μm pore size filters, aliquoted, and stored at -80°C until use.
For lentiviral transduction, $1x10^6$ target cells per well were seeded into 6-well plates and inoculated with lentivirus for 24 h. After washing, cells were cultured in antibiotics (puromycin or blasticidin) containing medium for selection.
To generate HepG2-NTCP ApoC1 KO cells, sgRNAs (sgRNA-1: 5'-

GGAGTTTGGAAACACACTGG-3’; sgRNA-2: 5’-TCCAAGGCACTGGAGACGTC-3’) were inserted into the LentiCRISPRv2 vector and confirmed by DNA sequence analysis. Stable ApoC1 knockout cells were selected via blasticidin resistance, cloned and subjected to western blot analysis to identify ApoC1 null clones.

##### *Immunoprecipitation*

After washing with cold TN buffer including 1x cOmplete EDTA-free Protease Inhibitor Cocktail (Roche, Switzerland), Pierce Protein A/G magnetic beads (Thermo Fisher, USA) were incubated with either antibodies of interest or IgG isotype controls for 30 mins at room temperature with constant rotation (10 μg antibody and 0.5 mg (50 μl) Protein A/G magnetic beads per sample), followed by one washing with cold TN buffer to remove uncoupled antibodies. Antibodies coated beads were resuspended in the sample (e.g. purified virus, supernatant or patient serum) and incubated for 3-4 h at 4°C on a rotating wheel. Beads were pelleted and washed with cold TN buffer four times. At the end, the beads were resuspended in 50 μl TN buffer for subsequent qPCR and western blot analyses.

##### *Immunogold labeling and electron microscopy*

Sucrose fractions containing HBV virions or fSVPs were diluted with TN buffer and applied to Amicon Ultra (30k) centrifugal filters to remove sucrose. HepAD38-derived samples were fixed with 2% paraformaldehyde (PFA) for 4 h at 4°C, while HepG2-HB2.7 samples were processed directly without prior fixation. Immunogold labelling was conducted as described previously[5]. In brief, 7 μl of different dilutions of sucrose-free HBV particles were applied to freshly glow-discharged carbon- and pioloform-coated 300-mesh copper grids (Science Services GmbH, Germany) for 5 mins at room temperature. After removing excess sample by touching with Whatman paper, grid-absorbed samples were washed with distilled water, incubated with blocking buffer (0.8% BSA, 0.1% fish skin gelatin, 50 mM glycine in PBS) for 20 mins, followed by incubation with primary antibodies for 20 mins at room temperature. After three times washing with PBS, the grids were incubated with Protein A-10 nm gold conjugates (Cell Microscopy Core, Netherlands) for 20 mins and washed three times with PBS, followed by two quick washes with distilled water. Grids were soaked into a droplet of 3% uranyl acetate solution and negatively stained with another 3% uranyl acetate droplet for 5 mins. The dye was removed using Whatman paper, and grids were air-dried at room temperature before imaging with a JEOL JEM-1400 transmission electron microscope (Jeol GmbH, Japan).

The antibodies used in this study are listed in Table S4.

***HBV preS1 binding assay***

The MyrB peptide (amino acid residues 2-48 of the HBV preS1 region with an N-terminal myristoylation) was synthesized by solid-phase synthesis[6]. Fluorophore-conjugated MyrB was generated by coupling the atto488-NHSester to the C-terminal lysine residue of the peptide ($MyrB^{atto}$). The cell binding assay of $MyrB^{atto}$ was performed as described previously[7]. In brief, cells grown on coverslips were washed with PBS and incubated with 200 nM peptide at 37°C for 30 mins, followed by washes with 2% BSA in PBS to remove unbound peptides. Cells were fixed with 4% PFA, stained with Hoechst dye for 5-10 mins at room temperature and mounted. Fluorescence images were captured using a Nikon Eclipse Ti Inverted Microscope and processed using the Image J software package.

***Statistical analysis***

The number of independent replicates is described in the figure legends and statistical analyses were performed using the GraphPad Prism v10 software package (GraphPad software, USA). Significance was determined using unpaired t test, one-way ANOVA or two-way ANOVA, depending on the experiment. P value lower than 0.05 were considered statistically significant, and stars in the graphs represent significance p values (*, $p < 0.05$, **, $p < 0.01$, ***, $p < 0.001$, ****, $p < 0.0001$).

#### Supplementary figures

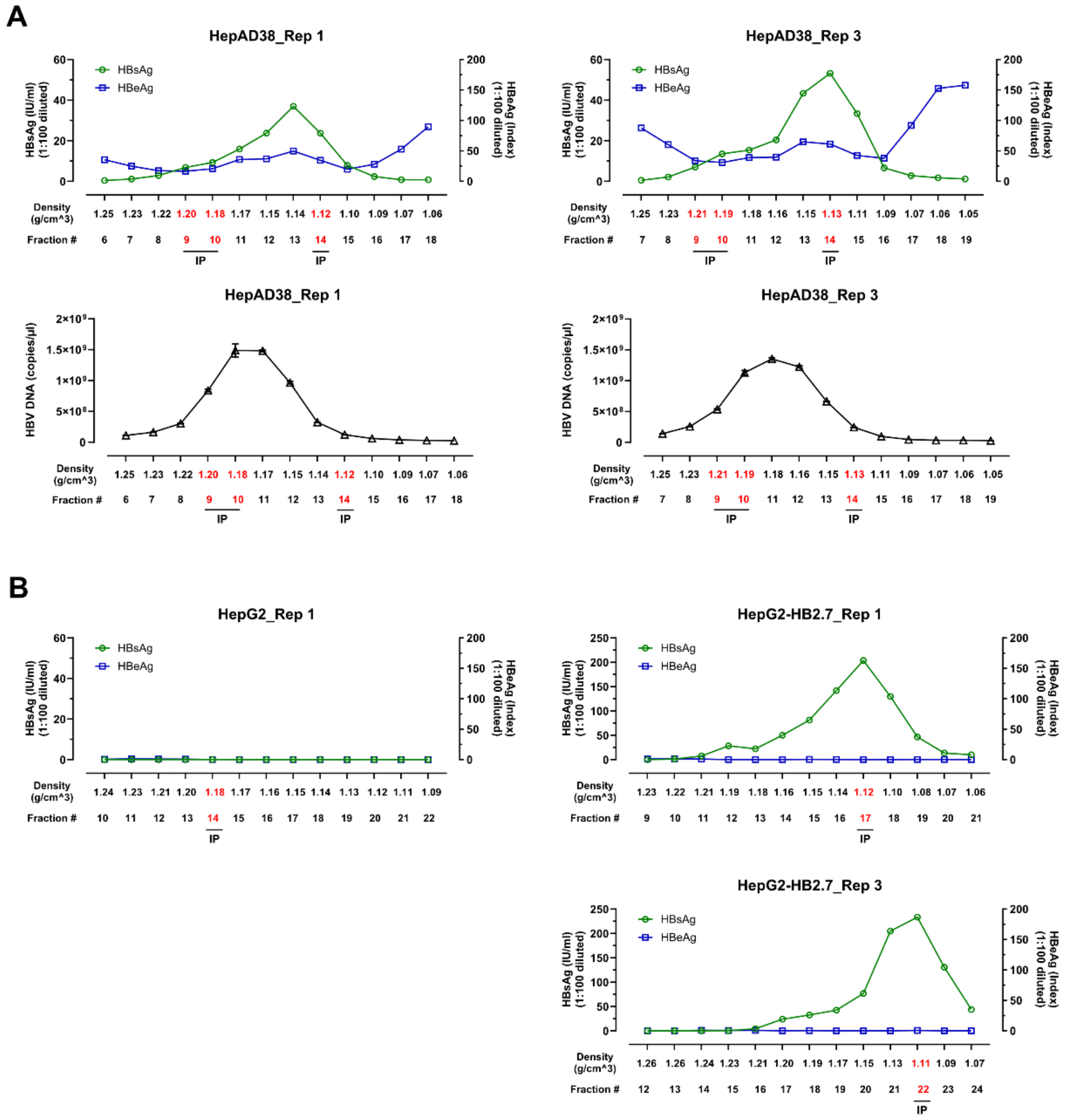

**Fig. S1. Characterization of sucrose gradient fractions.** Sucrose fractions of equilibrium density gradients were analyzed for HBsAg, HBeAg and HBV DNA levels by ELISA and qPCR, respectively. Cell lines from which the samples are derived and the number of the biological replicate are specified on the top of each panel. Sucrose densities of each fraction are specified below each lane. The red-highlighted fractions were used for HBsAg-specific pull-down.

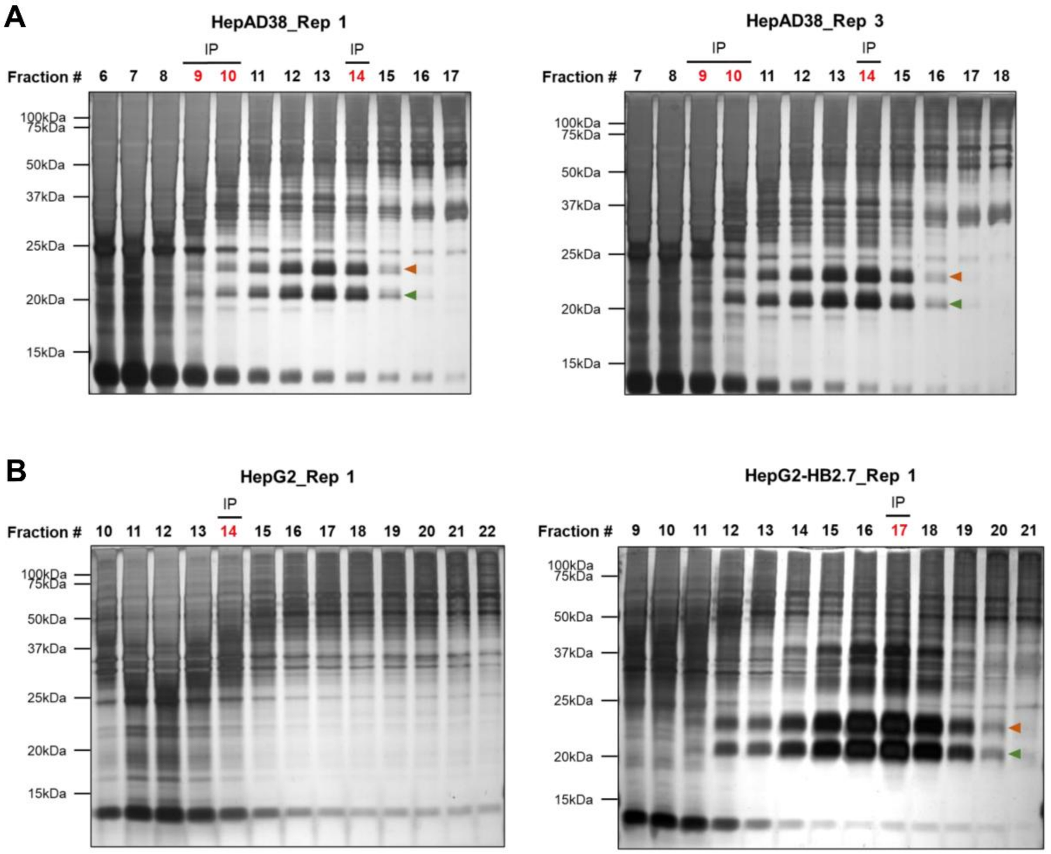

**Fig. S2. Total protein content in sucrose gradient fractions from control HepG2 and HBV particles-secreting cell lines.** Sucrose fractions of equilibrium density gradients were examined by silver gel staining to determine protein abundance and complexity in each sample. Cell lines from which the samples are derived and the number of the biological replicate are specified on the top of each panel. The red-highlighted fractions were used for HBsAg-specific immunoprecipitation. Arrowheads indicate glycosylated (orange) and non-glycosylated (green) small HBs proteins.

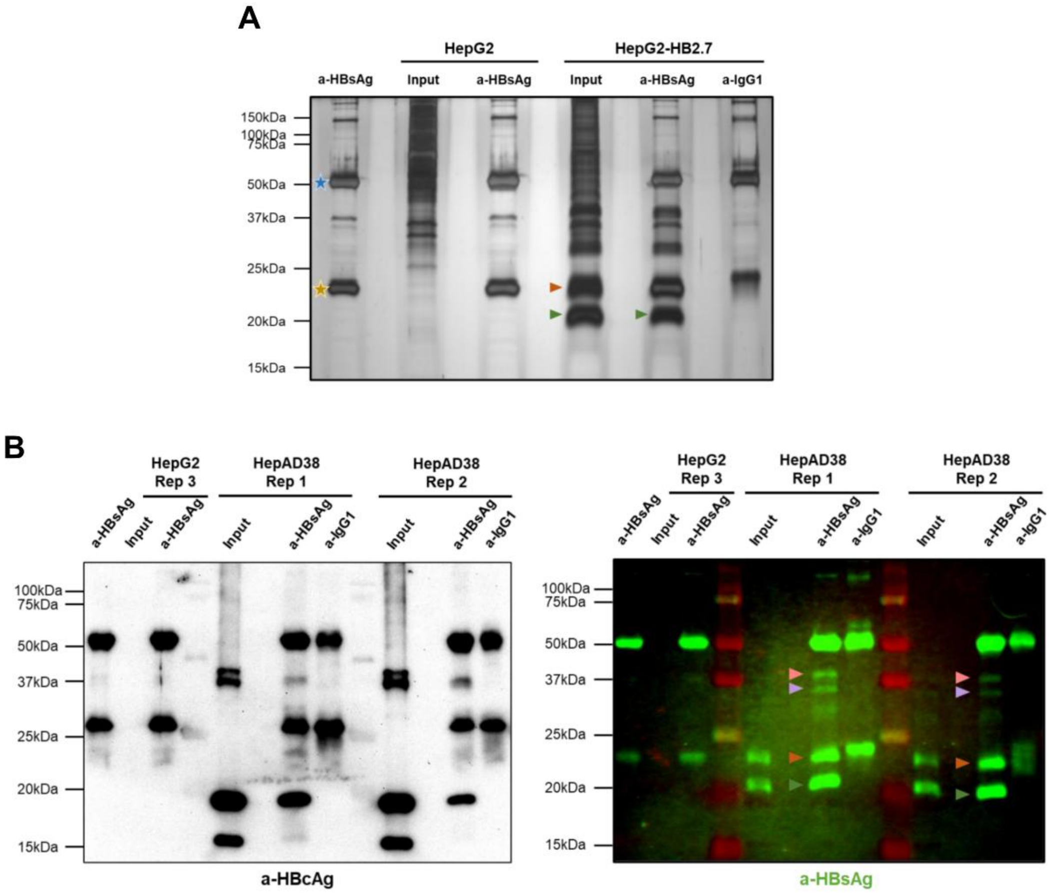

**Fig. S3. Immunoprecipitation and characterization of HBV subviral particles and virions.**

(A) Sucrose gradient fractions enriched for fSVPs that were isolated from HepG2-HB2.7 cells were subjected to immunoprecipitation with HBs-specific or IgG control antibodies and immunocomplexes were analyzed by SDS-PAGE and silver staining. Samples derived from naïve HepG2 cells were processed in parallel. Antibody-conjugated beads without input material were loaded as background control. Arrowheads indicate glycosylated (orange) and non-glycosylated (green) small HBs proteins; blue and gold stars indicate IgG heavy and light chains, respectively. (B) Left panel: Pull-down samples and input contained in the virion fractions, along with negative control samples from naïve HepG2 cells, were analyzed by HBcAg-specific western blot. Right panel: The blot was re-probed with an HBsAg-specific antibody. Arrowheads indicate glycosylated (orange) and non-glycosylated (green) small HBs proteins, as well as glycosylated (pink) and non-glycosylated (purple) large HBs proteins. Origin cell lines are specified on the top of each panel.

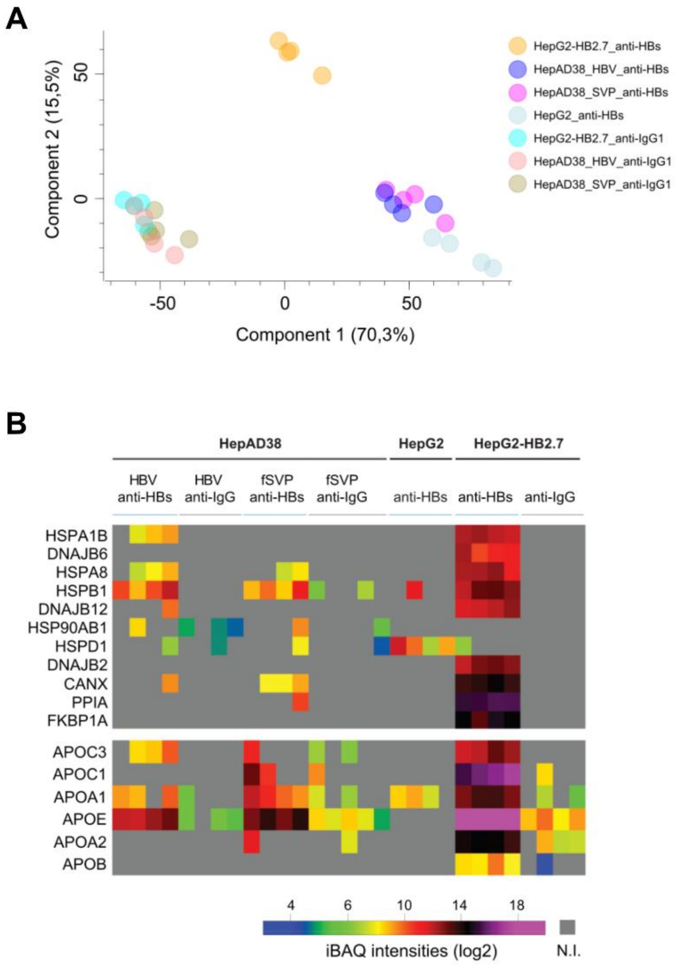

**Fig. S4. Refined analyses of proteome data.** (A) Principal component analysis (PCA) of biological replicates across HBV and filamentous SVP eluates after removal of outliers. (B) Functional enrichment analysis of the total proteins associated with purified HBV virions and fSVPs was performed using the online tool g:Profiler[8]. The analysis identified protein folding and plasma lipoprotein particle assembly groups, which are highlighted in the heatmap. Each column represents the proteome of a pull-down sample from one independent biological replicate. The color of the box represents the intensity-based quantification (iBAQ) of individual proteins (log2). N.I.=not identified.

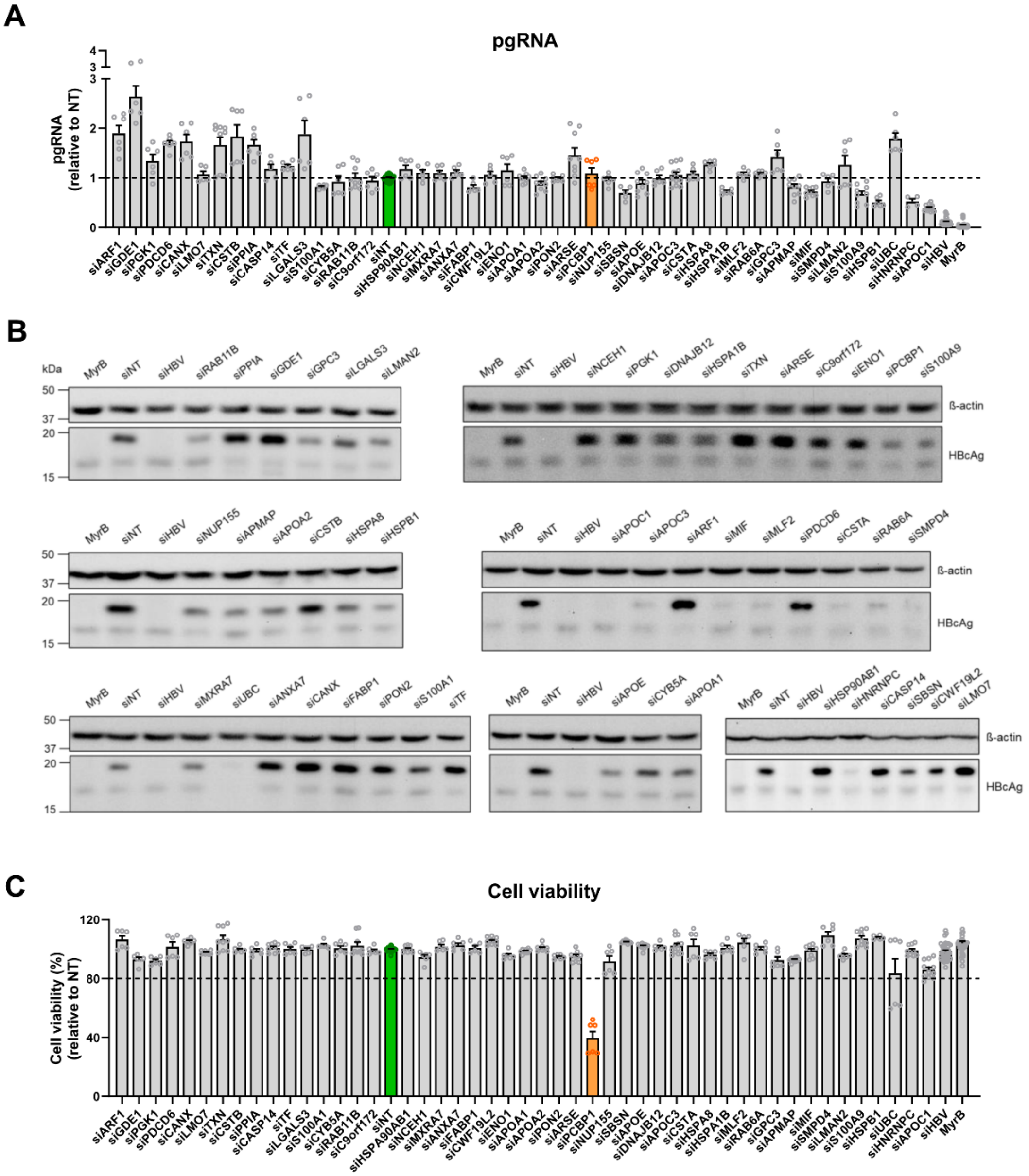

**Fig. S5. siRNA-based screen identifies novel HBV host-dependency factors.** (A) Pregenomic RNA (pgRNA) was quantified by RT-qPCR at 5 dpi. Values were normalized to siNT (set to 1; green bar). (B) Intracellular HBcAg was detected by western blot, and representative results are shown. (C) Cell viability was assessed at 5 dpi by WST-1 assay and values were normalized to siNT (set to 100%). The knock-down causing cytotoxicity (below 80% viability that we defined as our cut-off) is marked with the orange bar. Data are presented as mean ± SEM from at least two independent experiments performed in triplicate.

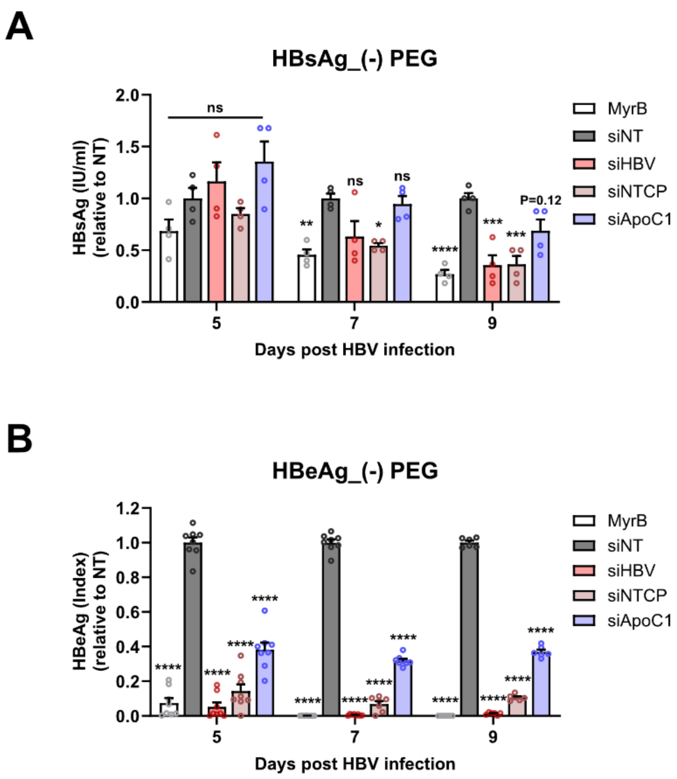

**Fig. S6.** (A-B) HepG2-NTCP A3 cells were transfected with siRNAs for 48 h and then infected with HBV with 1,200 GE/cell in the absence of 4% PEG8000 for 24 h. Secreted HBsAg (A) and HBeAg (B) were quantified by ELISA at 5, 7 and 9 dpi. Data are presented as mean ± SEM from three independent experiments. Two-way ANOVA was applied; *p<0.05, **p<0.01, ***p<0.001, ****p<0.0001, ns-not significant (p>0.05).

MyrB, cells treated with the entry inhibitor Bulevirtide.

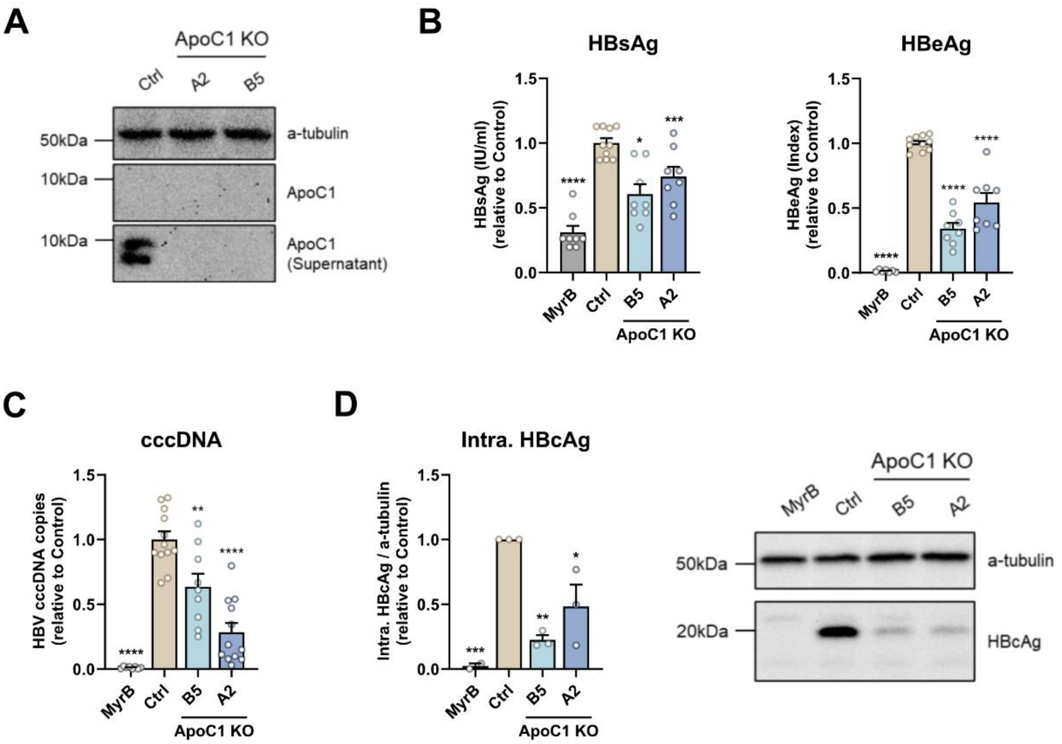

**Fig. S7. ApoC1 knockout attenuates HBV infection.** HepG2-NTCP A3 cells were transduced with a lentivirus (LentiCRISPRv2-Blast/ApoC1-sgRNA) encoding Cas9 and a ApoC1-specific sgRNA. Upon selection with blasticidin, stable cell clones were isolated and expanded. For control purposes, HepG2-NTCP A3 cells were transduced with a lentivirus encoding Cas9 and a non-targeting sgRNA. (A) ApoC1 loss in the stable cell clones was determined by western blot. (B-D) ApoC1 knockout (KO) cell clones were infected with HBV (500 GE/cell) for 24 h in parallel to HepG2-NTCP A3 cells containing the non-targeting sgRNA (Ctrl). Cells treated with the entry inhibitor Bulevirtide (MyrB) served as positive control. (B) Secreted HBsAg and HBeAg were measured by ELISA at 7 dpi. (C) HBV cccDNA was determined by qPCR at 7 dpi. (D) Intracellular HBcAg was detected by western blot at 7 dpi and quantified using Image Lab. Values were normalized to alpha-tubulin serving as loading control. One representative WB result is shown. Data are presented as mean ± SEM from at least three independent experiments. B, C, D, One-way ANOVA; *p<0.05, **p<0.01, ***p<0.001, ****p<0.0001.

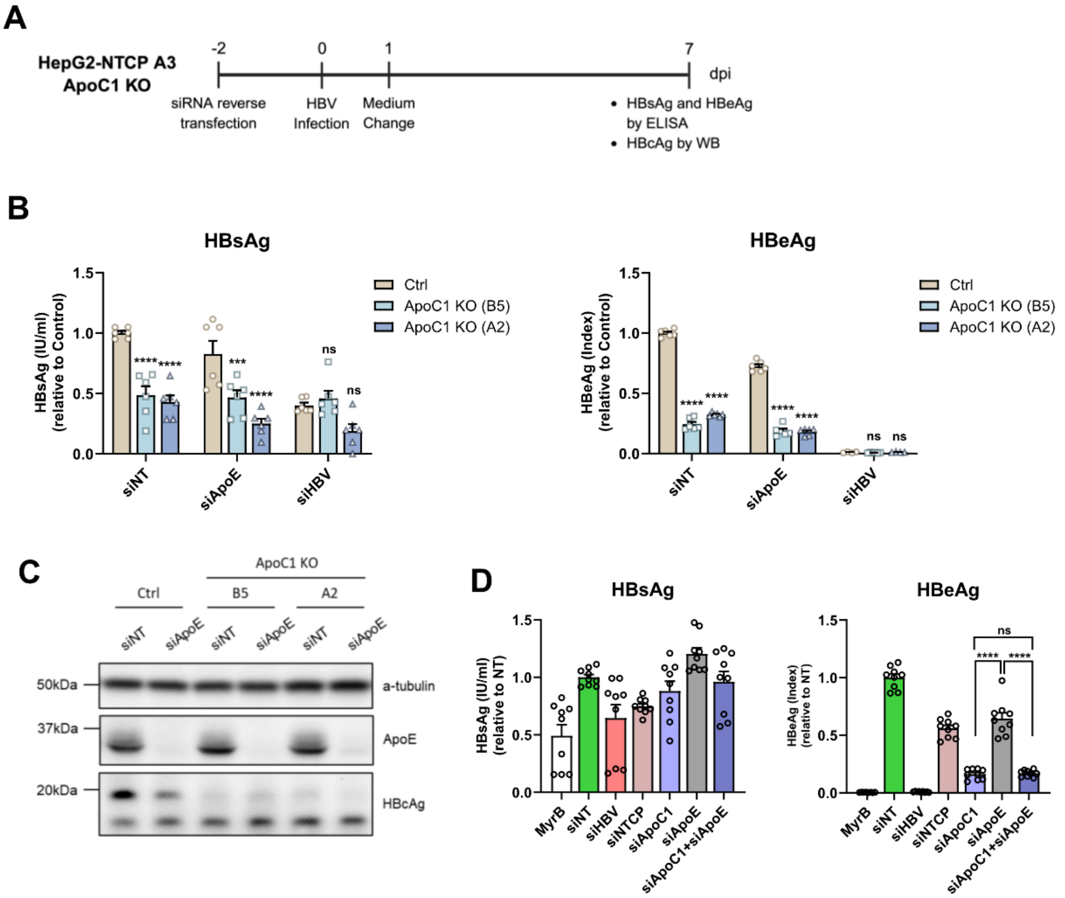

**Fig. S8. Impact of ApoE knockdown in ApoC1 deficient cells on HBV.** (A) HepG2-NTCP A3 ApoC1 KO cell clones or HepG2-NTCP A3 cells containing the non-targeting control sgRNA (Ctrl) were transfected with ApoE-targeting siRNAs for 48 h and subsequently infected with HBV. After removal of the inoculum, cells were cultured for an additional 7 days. Cells transfected with the HBV-specific siRNA served as positive control. (B) Secreted HBsAg and HBeAg were measured by ELISA at 7 dpi. (C) Intracellular HBcAg and ApoE were detected by western blot; one representative result is shown. (D) SiRNA-mediated single and double knockdown of ApoC1 and ApoE were performed in HepG2-NTCP A3 cells, and secreted HBsAg and HBeAg were measured by ELISA at 5 dpi. Data are presented as mean ± SEM from three independent experiments. B, Two-way ANOVA; D, One-way ANOVA; ***$p<0.001$, ****$p<0.0001$, ns-not significant ($p>0.05$).

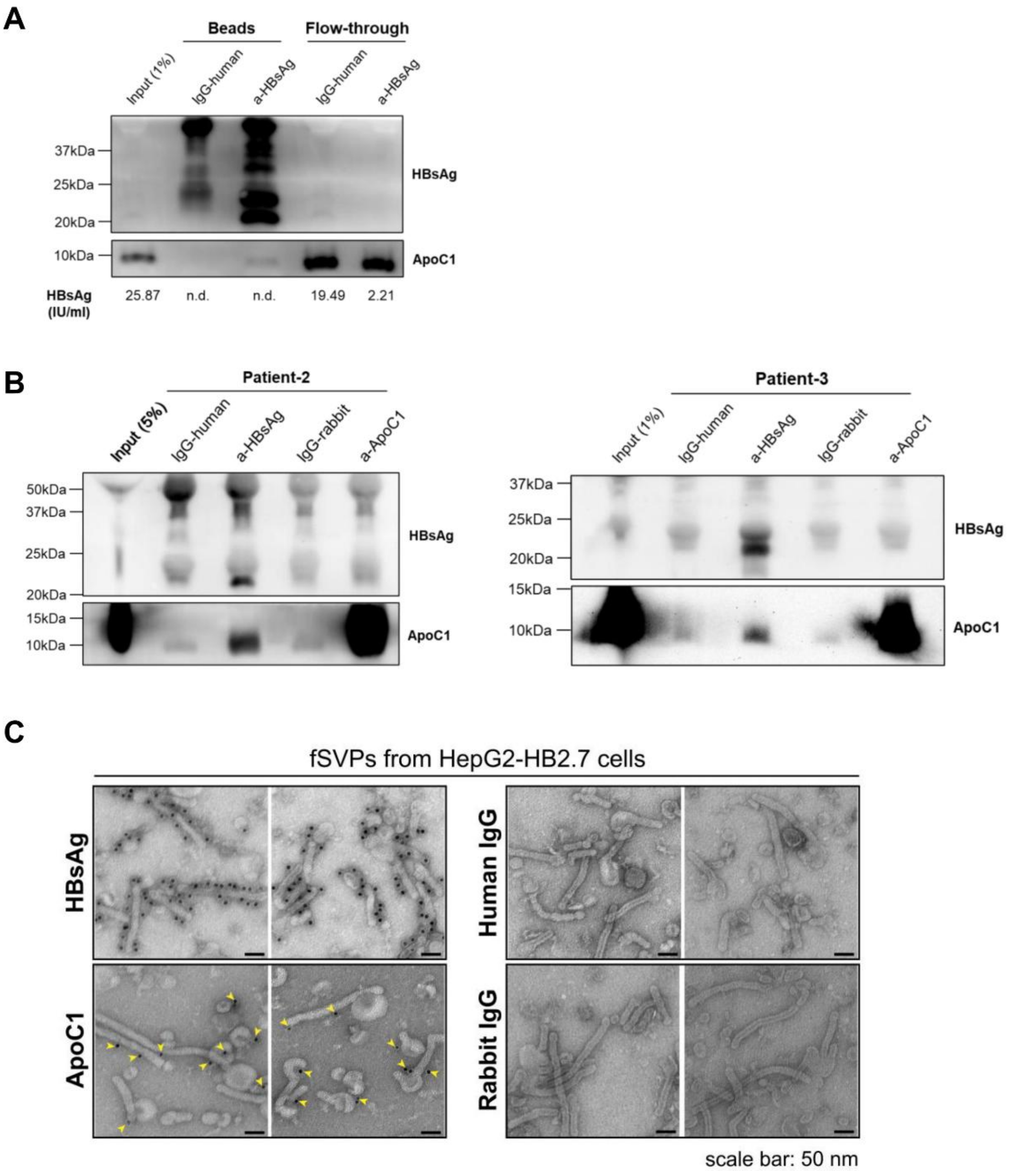

**Fig. S9. ApoC1 associates with filamentous SVPs.** (A) Huh7 cells were transfected with a plasmid containing a 1.1-overlength HBV genome. Supernatants collected from day 4-7 were used for immunoprecipitation with HBsAg-specific and IgG control antibodies, the latter from matching species. HBsAg ELISA values from input and flow-through fractions are given below each lane. N.d., non-detectable. (B) HBsAg-specific immunoprecipitation of HBV particles contained in sera of two patients with CHB. Samples were analyzed by western blot. (C) fSVPs released from HepG2-HB2.7 cells and purified by heparin affinity chromatography, size exclusion chromatography and sucrose gradient centrifugation were immunogold labeled for HBsAg and ApoC1 and visualized by transmission electron microscopy. Species-matching IgG antibodies served as specificity control. For each condition, two representative view fields are shown. ApoC1-specific immunogold is marked with yellow arrow heads. Scale bar: 50nm.

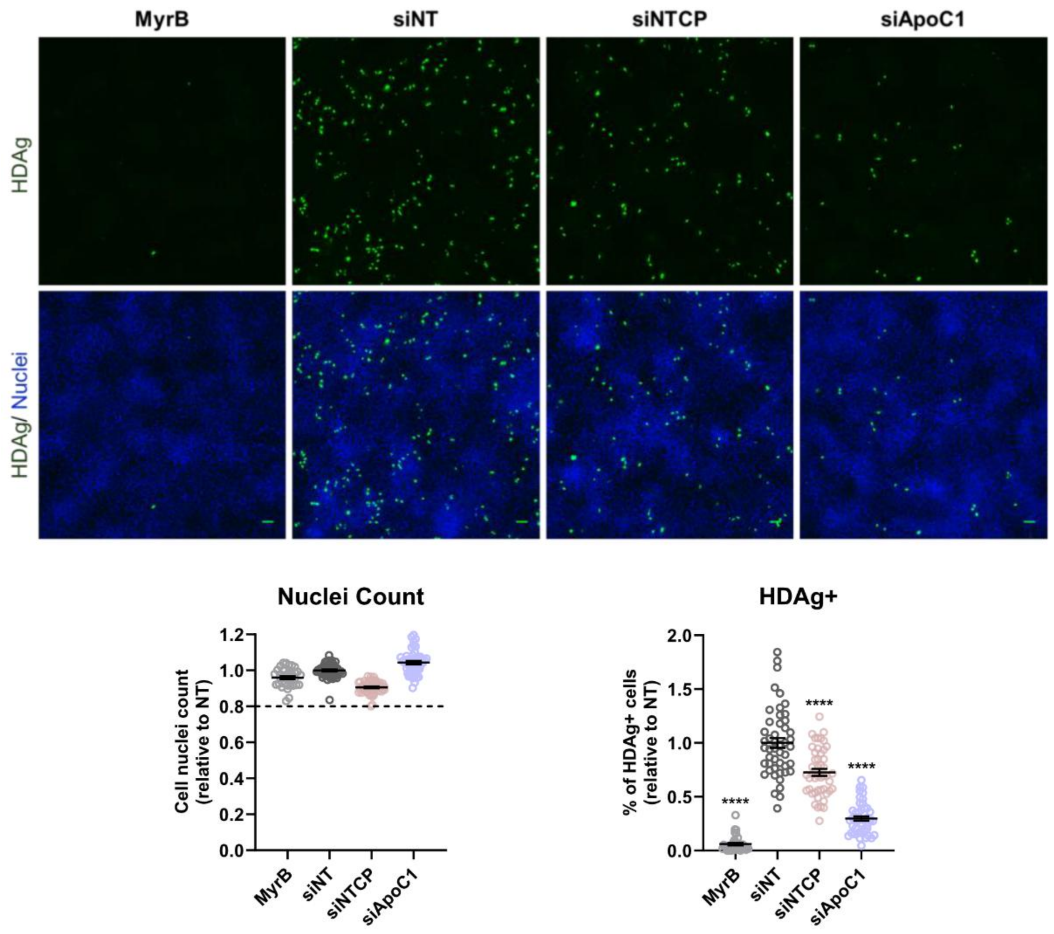

**Fig. S10. ApoC1 knockdown reduces HDV infection/replication.** Huh7-NTCP cells were reverse transfected with specified siRNAs for 48 h and subsequently infected with HDV at 5 GE/cell in the presence of 4% PEG8000 for 24 h. Intracellular HDAg was assessed by immunofluorescence at 5 dpi. Green and blue signals on the upper panel depict the staining of HDAg and nuclei, respectively. Quantifications of nuclei number and HDAg positive cells are shown on the lower panel. Cells treated with the entry inhibitor Bulevirtide (MyrB) served as positive control. Scale bar: 100 µm. Data are presented as mean ± SEM from three independent experiments; One-way ANOVA analysis. ****$p<0.0001$.

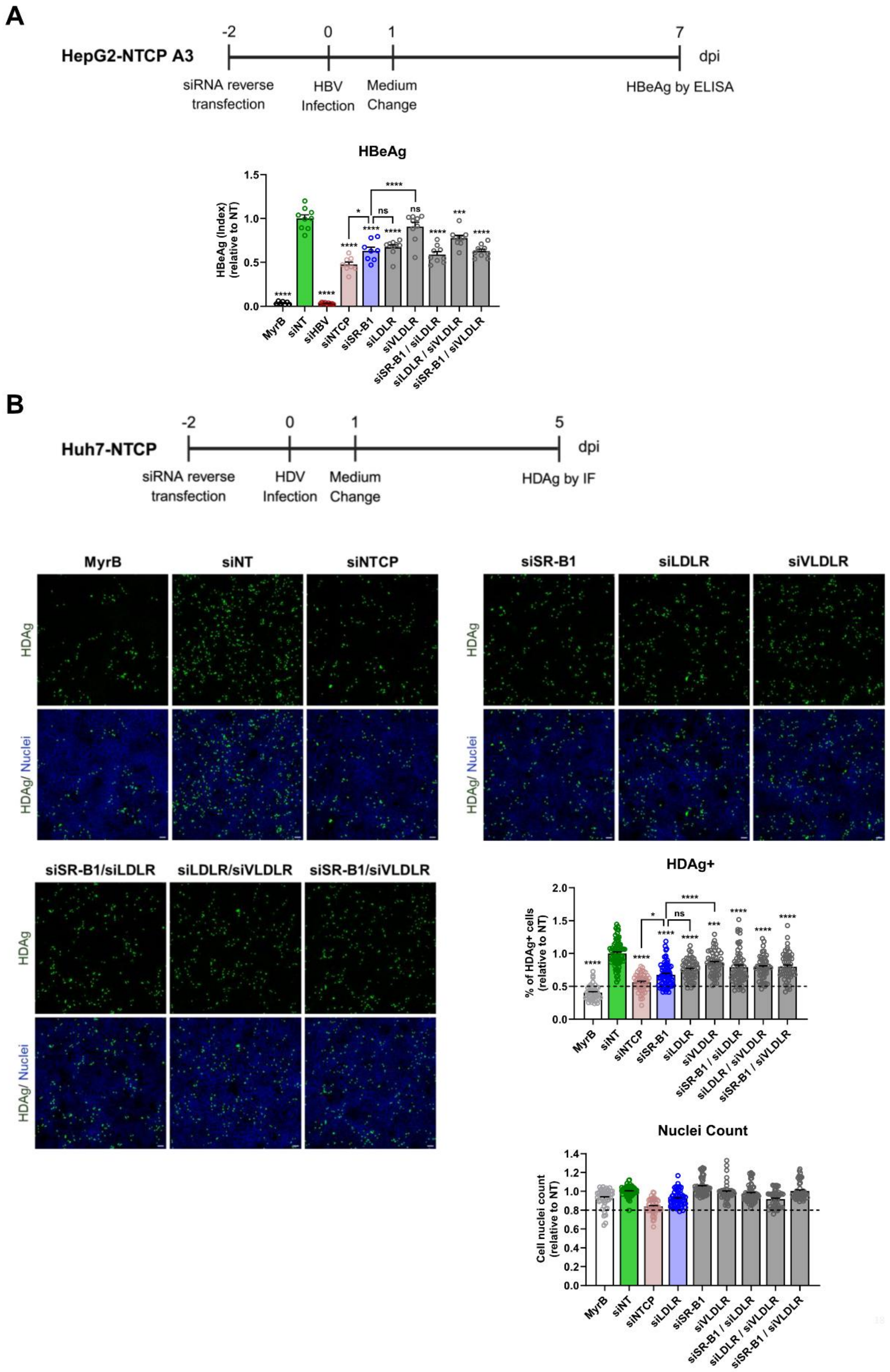

**Fig. S11. Role of lipoprotein receptors in HBV and HDV infection.** (A) HepG2-NTCP A3 or (B) Huh7-NTCP cells were reverse transfected with siRNA targeting lipoprotein receptors specified on the bottom for 48 h and subsequently infected with HBV or HDV, respectively. The experiments were performed under FBS-free conditions during infection. (A) Secreted HBeAg was measured by ELISA at 7 dpi. (B) Intracellular HDAg was assessed by immunofluorescence at 5 dpi. Green and blue signals depict the staining of HDAg and nuclei, respectively. Quantifications of nuclei number and HDAg positive cells are shown. Scale bar: 100μm. Data are presented as mean ± SEM from three independent experiments with One-way ANOVA analysis. *$p<0.05$, ***$p<0.001$, ****$p<0.0001$, ns-not significant ($p>0.05$).

#### Supplementary tables

**Table S1. Intensity values of viral peptides identified by AP-LC-MS/MS.** List of viral peptide sequences and iBAQ intensities across all samples of 5 biological replicates. Missing numerical values in the dataset were represented as NaN (Not a Number).

**Table S2. Intensity values underlying proteomic analysis of filamentous SVPs, HBV virions and related controls.** (A) List of all unnormalized iBAQ intensities across all biological replicates and experimental groups. No outliers or contaminants removed. (B) List of median-normalized iBAQ intensities across samples and experimental groups. Common contaminants and outliers were removed, and iBAQ intensity values normalized by the median of each sample, and shifted of 12 Log2 values into positive numerical space. Proteins curated as HBV and filamentous SVP specific (proteins identified in at least 3 out of 4 biological replicates and enriched at least 8-fold compared to the corresponding IgG control) are indicated (n=168 total). Additionally, proteins belonging to the GO terms "protein folding" and "plasma lipoprotein particle assembly" as well as proteins selected for siRNA screening are indicated. Missing numerical values in the dataset were represented as NaN (Not a Number).

**Table S3. Sequences of primers used for qPCR and RT-qPCR.**

| Primer | Sequence (5'-3') |
|---|---|
| HBV DNA | F: GTTGCCCGTTTGTCCTCTAATTC |
| | R: GGAGGGATACATAGAGGTTCCTTGA |
| HBV total RNA | F: TCAGCAATGTCAACGACCGA |
| | R: TGCGCAGACCAATTTATGCC |
| pgRNA | F: CTCCTCCAGCTTATAGACC |
| | R: GTGAGTGGGCCTACAAA |
| HDV RNA | F: GCGCCGGCTGGGCAAC |
| | R: TTCCTCTTCGGGTCGGCATG |
| HDV RNA probe | FAM- CGCGGTCCGACCTGGGCATCCG-BHQ |
| APOA1 | F: CCCTGGGATCGAGTGAAGGA |
| | R: CTGGGACACATAGTCTCTGCC |
| APOA2 | F: CTGTGCTACTCCTCACCATCT |
| | R: CTCTCCACACATGGCTCCTTT |
| APOC1 | F: TCCAGTGCCTTGGATAAGCTG |
| | R: GGCTGATGAGTTCCCGAGC |
| APOC3 | F: TTACATGAAGCACGCCAC |
| | R: CTCCAGTAGTCTTTCAGGGAAC |
| APOE | F: GTTGCTGGTCACATTCCTGG |
| | R: GCAGGTAATCCCAAAAGCGAC |
| GAPDH | F: TCGGAGTCAACGGATTTGGT |
| | R: TTCCCGTTCTCAGCCTTGAC |
| ß-globin | F: CAGGTACGGCTGTCATCACTTAGA |
| | R: CATGGTGTCTGTTTGAGGTTGCTA |
| cccDNA | F: GTGGTTATCCTGCGTTGAT |
| | R: GAGCTGAGGCGGTATCT |
| cccDNA probe | FAM-AGTTGGCGAGAAAGTGAAAGCCTGC-TAMRA |

F, forward primer; R, reverse primer

#### Table S4. Antibodies used for western blot and immunostaining.

| Antibody | Supplier | Catalog No. | Host | Clonality | Dilution |
|---|---|---|---|---|---|
| a-tubulin | Sigma-Aldrich | T5168 | Mouse | Monoclonal | 1:8000 (Western blot) |
| ß-actin | Sigma-Aldrich | A5441 | Mouse | Monoclonal | 1:5000 (Western blot) |
| HBcAg | Dako | B0586 | Rabbit | Polyclonal | 1:3000 (Immunofluorescence) |
| HBcAg (H746) | Christa Kuhn | N/A | Rabbit | Polyclonal | 1:2000 (Western blot) |
| HBsAg (HBC34) | Humabs Biomed | N/A (gift from Davide Cortin) | Human | Monoclonal | 1:3000 (Immunofluorescence) |
| HBsAg (HBD87) | Humabs Biomed | N/A (gift from Davide Cortin) | Human | Monoclonal | 1:5000 (Western blot) |
| HDAg (FD3A7) | Absolute antibody | Ab03320-23.0 | Rabbit | Monoclonal | 1:3000 (Immunofluorescence) |
| ApoC1 | abcam | ab198288 | Rabbit | Monoclonal | 1:1000 (Western blot)<br>1:100 (Immunogold labelling) |
| ApoE | Sigma-Aldrich | AB947 | Goat | Polyclonal | 1:5000 (Western blot) |
| Normal Rabbit IgG | Cell Signaling Technology | 2729 | Rabbit | Polyclonal | Immunoprecipitation |
| Human IgG1 Isotype Control | Novus Biologicals | DDXCH01P-100 | Human | Polyclonal | Immunoprecipitation |
| Anti-Mouse IgG HRP | Sigma-Aldrich | A4416-5X1ML | Goat | Polyclonal | 1:10000 (Western blot) |
| Anti-Rabbit IgG HRP | Sigma-Aldrich | A6154-5X1ML | Goat | Polyclonal | 1:20000 (Western blot) |
| Anti-Goat IgG HRP | Santa Cruz | sc-2020 | Donkey | Polyclonal | 1:2000 (Western blot) |
| Anti-Human IgG H&L HRP | abcam | ab97165 | Goat | Polyclonal | 1:10000 (Western blot) |
| Goat anti-human IgG (H+L), Alexa Fluor 488 | Invitrogen | A11013 | Goat | Polyclonal | 1:1000 (Immunofluorescence) |
| Goat anti-rabbit IgG (H+L), Alexa | Invitrogen | A11008 | Goat | Polyclonal | 1:1000 (Immunofluorescence) |

| | | | | | |
|---|---|---|---|---|---|
| Fluor 488 | | | | | |
| Donkey anti-rabbit IgG (H+L), Alexa Fluor 647 | Invitrogen | A31573 | Donkey | Polyclonal | 1:1000 (Immunofluorescence) |
